# Deciphering the RNA-based regulation mechanism of the phage-encoded AbiF system in *Clostridioides difficile*

**DOI:** 10.1101/2025.04.15.648962

**Authors:** Marion Saunier, Adeline Humbert, Victor Kreis, Johann Peltier, Arianna Tisba, Sylvie Auxilien, Marion Blum, Isabelle Caldelari, Jean-François Lucier, Joe Ueda, Daniel Gautheret, Claire Toffano-Nioche, Jessica Andreani, Louis-Charles Fortier, Olga Soutourina

**Affiliations:** Université Paris-Saclay, CEA, CNRS, Institute for Integrative Biology of the Cell (I2BC), 91198, Gif-sur-Yvette, France; Architecture et Réactivité de l’ARN, UPR9002, CNRS, Université de Strasbourg, Strasbourg, France; Biology Department, Université de Sherbrooke, Sherbrooke, QC J1K 2R1, Canada; Department of Microbiology and Infectious Diseases, Université de Sherbrooke, Sherbrooke, QC J1E 4K8, Canada

**Keywords:** *Clostridioides difficile*, Abi-like system, type III toxin-antitoxin, RNA antitoxin, RNA-protein interaction, prophage, hypervirulent strain

## Abstract

*Clostridioides difficile* is the major cause of nosocomial infections associated with antibiotic therapy. The severity of *C. difficile* infections increased worldwide with the emergence of hypervirulent strains, including 027 ribotype epidemic strains. Many aspects of *C. difficile*’s adaptation strategies during pathogenesis remain poorly understood. This pathogen thrives in gut communities that are rich in microbes and phages. To regulate horizontal transfer of genetic material during its infection cycle, *C. difficile* relies on diverse mechanisms. More specifically, CRISPR (clustered regularly interspaced short palindromic repeats)-Cas and Toxin-Antitoxin (TA) systems contribute to prophage maintenance, prevention of phage infection, and stress response. Abortive infection (Abi) systems can provide additional lines of anti-phage defense. RNAs have emerged as key components of these systems including CRISPR RNAs and antitoxin RNAs within type I and type III TA. We report here the identification of a new AbiF-like system within a prophage of the hypervirulent *C. difficile* strain R20291. It is associated with an Abi_2/AbiD/F protein family largely distributed in Bacillota and Pseudomonadota with structural links to ancestral Cas13 proteins at the origin of the RNA-targeting CRISPR-Cas13 systems. We demonstrated toxic activity of the AbiF*_Cd_* protein in *C. difficile* and in *Escherichia coli* and negative regulation of the *abiF_Cd_* expression by an associated non-coding RNA RCd22. RCd22 contains two conserved *abiF* motifs and is active both in *cis* and *in trans* to neutralize the toxin by direct RNA-protein interaction, similar to RNA antitoxin in type III TA. A mass spectrometry interactomics analysis of protein fractions from MS2-Affinity Purification coupled with RNA sequencing (MAPS) revealed the AbiF*_Cd_* protein among the most enriched RCd22 partners in *C. difficile*. Structural modeling of the RNA-protein complex and mutagenesis analysis revealed key positions on both protein and RNA partners for this interaction and toxic activity. In summary, these findings provide valuable insights into the mechanisms of interaction between bacteria and phages, which are pertinent to the advancement of phage therapy, genome editing, epidemiological surveillance, and the formulation of novel therapeutic approaches.

## Introduction

Bacterial evolution has been profoundly affected by viruses, namely bacteriophages (or phages). To defend themselves against phages, bacteria have developed efficient anti-phage defense systems such as Clustered Regularly Interspaced Short Palindromic Repeats (CRISPR), CRISPR-associated (Cas), Toxin-Antitoxin (TA), Restriction-Modification (RM), superinfection exclusion (Sie) and Abortive Infection (Abi) systems. Abi is a common defense mechanism present in most bacteria. Upon activation, Abi induces cell death (or dormancy) before the infecting phage can complete its replication cycle, thus protecting the colony from phage spreading (1). This defense strategy is associated with numerous newly discovered immune systems (2). The total number of defense systems described so far exceeds 130, revealing the complexity of bacterial immunity and the diversity of associated mechanisms (3).

*Clostridioides difficile,* a human pathogen responsible for antibiotic-associated infections, evolves in a phage-rich environment inside the intestinal tract. To cope with the presence of phages and other invaders, this bacterium has developed several defense mechanisms. In addition to a well-characterized type I-B CRISPR-Cas system (4–7), other anti-phage systems were identified such as RM (8) and superinfection exclusion (9). A putative *abiF-*like gene was also found in the *C. difficile* phi027 prophage (9), φC2 (10), and φCD2301 (11) phage genomes, but its function and contribution to the defense mechanisms remain uncharacterized. The AbiF system was originally described in *Lactococcus lactis* (12). It shares structural similarities with AbiD and AbiD1, two other systems of *L. lactis,* giving a single AbiD/F group of abortive infection systems (13). This group belongs to the superfamily Abi_2, widely distributed among bacteria. In *L. lactis*, AbiF provides resistance to phage infection by interfering with DNA replication (12) while AbiD1 interferes with a phage-encoded endonuclease (14,15). Aside from what is known in *L. lactis*, the function and activity of this group of Abi systems remain poorly characterized in other species, and the exact mechanism of action for both AbiD1 and AbiF proteins remains to be defined.

Given the toxicity of Abi systems for the bacterial cell itself, they must be tightly regulated. Most Abi systems are composed of a sensing module responding to phage infection, and an effector module triggering cell death or dormancy (1,2). The sensing module usually detects a conserved phage protein directly, or sometimes indirectly through the effect of phage infection on the host, like inhibition of the cell machinery (2). In *L. lactis* AbiD1, a structured 5’UTR region on the *abiD1* mRNA was shown to contribute to the control of *abiD1* translation (15). During phage infection, a conserved phage protein binds to the 5ʹUTR of the *abiD1* mRNA, leading to the activation of *abiD1* translation (15). A short premature terminated transcript has been detected upstream of the *abiD1* coding sequence, suggesting that the AbiD1 system could be also regulated at the transcriptional level (16). Another example of an RNA-based mechanism associated with the abortive infection strategy is the type III toxin-antitoxin (TA) system. In this case, the antitoxin is a non-coding RNA (ncRNA) that neutralizes the toxin protein through an RNA-protein interaction (17,18). This system is activated following the inhibition of host transcription during phage infection, leading to the release of the stable toxin (19).

ncRNAs are key components of various regulatory mechanisms that control virulence in major pathogens. They also contribute to anti-phage systems including CRISPR-Cas, TA and recently discovered defense mechanisms such as CBASS (20) and retrons (21,22), both considered as Abi systems. Numerous ncRNAs were identified in *C. difficile*, including CRISPR RNA, *cis* and *trans* riboregulators and riboswitches, showing the importance of RNA-dependent regulation mechanisms in this human pathogen (4). Our previous work demonstrated that CRISPR arrays and type I TA modules are frequently associated with prophages in *C. difficile*. They provide efficient defense against other invading phages, and/or contribute to the stability of these regions (6,7,23). In this study, we describe RCd22, a ncRNA specific to the hypervirulent *C. difficile* strain R20291. Its sequence was found upstream of a putative *abiF-*like gene located in the lysis module of phi027, a prophage highly conserved among ribotype 027 *C. difficile* isolates (24,25). This work provides the characterization of a putative AbiF-like system in *C. difficile*, and reveals the tight regulation of this system by a ncRNA.

## Results

### Identification of an AbiF-like system conserved in *C. difficile* hypervirulent ribotype 027 strains

Abi systems have never been functionally characterized in clostridial species, although bioinformatics analyses revealed the presence of putative AbiF systems in prophages of *C. difficile* (9–11) (Fig. 1). In this study, we sought to characterize the function of a putative AbiF system located in the phi027 prophage present in the chromosome of the hypervirulent strain R20291 (9) (Fig. 1A) and generally well-conserved in ribotype 027 isolates of *C. difficile* (24,25). This putative AbiF system is encoded by a single gene *CDR20291_1462* (GenBank accession number: CBE04038.1), hereafter named *abiF_Cd_*. A BlastP search using this protein sequence as the query revealed homologs in the genomes of *C. difficile* phage φC2 (NCBI accession number: YP_001110753.1, 99.88% identity) (5,10) and φCD2301 (NCBI accession number: QVW56672.1, 99.54% identity) (Fig. 1B) (11). The hypothetical AbiF_Cd_ protein belongs to the superfamily Abi_2 (cl01988), containing the Abi_2 conserved domain (pfam07751) and the AbiF conserved domain (COG4823). The AbiF protein originally identified in a *L. lactis* plasmid shares 34.39% identity with AbiF_Cd_. Blast analyses of Abi_2/AbiF conserved domains were done on the complete genomes of 46,864 bacterial and 681 archaeal species. Results showed that this protein family is largely distributed in bacteria belonging to the group of Bacillota, comprising Bacilli and Clostridia classes. It is also found in the group of Pseudomonadota, comprising alpha, beta and gammaproteobacteria classes (Fig. 2A). The phylogenetic tree also showed that the Rickettsiales order (alphaproteobacteria class), *Borreliaceae* family (betaproteobacteria class) and several other genera and species lack the *abiF/abi_2* gene. Interestingly, when the *abiF/abi_2* gene was detected in a species, it was not necessarily detected in all genomes of that species. For example, among 1,800 genomes of *Staphylococcus aureus* analyzed, 200 contain no *abiF/abi_2* homologous genes (11%). Among 170 genomes of *C. difficile*, 106 lack the *abiF/abi_2* genes (62%). Bacterial species generally carry between one and up to eight copies of a putative *abi_2*/*abiF* gene in their genome (red bars in Fig. 2A), with a majority carrying one or two copies per genome. In *C. difficile* strains carrying a putative *abiF* gene, only one copy is present. The biological function of the AbiF system originally found on a *L. lactis* plasmid was characterized (12), but the function of the chromosome-encoded version remains unknown. Analysis of the gene environment and genome organization could give clues regarding the biological function of the putative AbiF/Abi_2 protein. We therefore sought for genes located near *abiF/abi_2* in the 7,828 annotated genomes carrying at least one *abiF/abi_2* gene homolog. We evaluated the annotated functions of gene products five positions upstream and five positions downstream from the *abiF/abi_2* gene (Fig. 2B). We grouped gene products according to their biological function. Genes present with an abundance of less than 10% were grouped in the category “others”. Despite the functional variability in the closest genomic environment, defense mechanisms and mobilome-associated gene functions were enriched in the proximity of the *abiF/abi_2* gene at the positions - 1 and +1. The presence of *abiF/abi_2* in prophages or transposons could explain the large distribution of this gene in bacteria. Other copies of *abiF/abi_2* (assigned to the defense mechanism functional class) were frequently found at the −1 or +1 position, suggesting potential gene duplication events. Other gene associations implying translation and posttranslational modification were found downstream from the *abiF/abi_2* gene, while genes related to nucleotide metabolism and transcription were found upstream of *abiF/abi_2*. Of note, all putative *abiF* genes in *C. difficile* strains are located between a predicted phage holin and a gene encoding an endolysin, within the lysis module of a prophage genome (Fig. 1B). The association with lysis module was also observed in other bacterial species, but the *abiF/abi_2* gene was located downstream from this module. Therefore, the location of *abiF* inside the lysis module is specific to *C. difficile* strains (Fig. 1A). The location of a putative anti-phage system inside a prophage is intriguing. It could provide benefits for both the bacterial host and the prophage, by protecting bacteria from infections by unrelated phages, and/or stabilizing the prophage in the bacterial chromosome, as shown for type I toxin-antitoxin systems (23,26,27). We further investigated the colocalization of the *abiF/abi_2* gene with prophage elements. Of the 7,828 annotated genomes carrying at least one *abiF/abi_2* gene homolog, 2,600 representative genomes were analyzed with geNomad (28). The coordinates of the identified prophages were then compared with the coordinates of the *abiF/abi_2* gene to detect overlaps. A total of 288 prophages were found to carry an *abiF/abi_2* gene homolog. Analysis of the −5 to +5 genomic environment surrounding *abiF/abi_2* revealed that the category “Mobilome: prophages, transposons” was enriched in the −1 and +1 position, as well as “cell wall/membrane/envelope biogenesis” (Fig. S1A). The latter category is consistent with *abiF/abi_2* being closely associated with the lysis module in prophages (i.e. holin and endolysin genes). In the 2,312 genomes where *abiF/abi_2* was not located on a prophage, the functions associated with “Defense mechanisms” and “Replication, recombination and repair” were enriched (Fig. S1B).

**Fig. 1.**
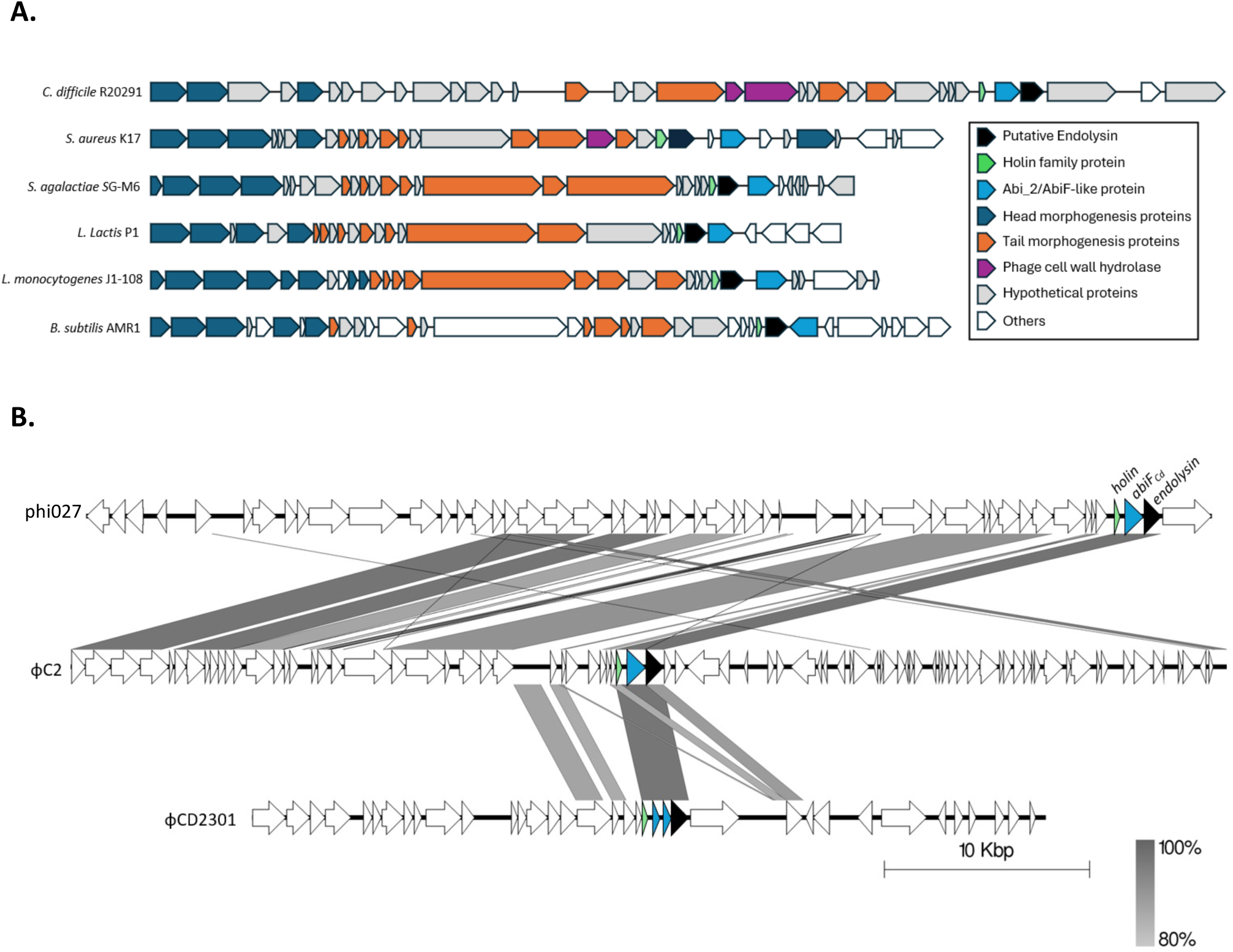
- Identification of putative AbiF/Abi_2 systems associated with phage lysis module. (**A**) Localization of putative *abiF/abi_2* genes in selected bacterial genomes. (**B**) Comparison of the genome sequence of three *C. difficile* prophages: phi-027 prophage (top), φC2 (middle) and φCD2301 (bottom). Predicted ORFs and the direction of transcription are indicated by arrows. Green arrows represent putative *holin* genes; dark blue arrows represent putative *abiF*-like genes; black arrows represent putative *endolysin* genes. Conserved regions are shaded in grey and color intensity corresponds to sequence identity level (80% to 100%). Genomic comparisons were performed using BLASTn and the figure was produced using Easyfig 2.2.5.

**Fig. 2.**
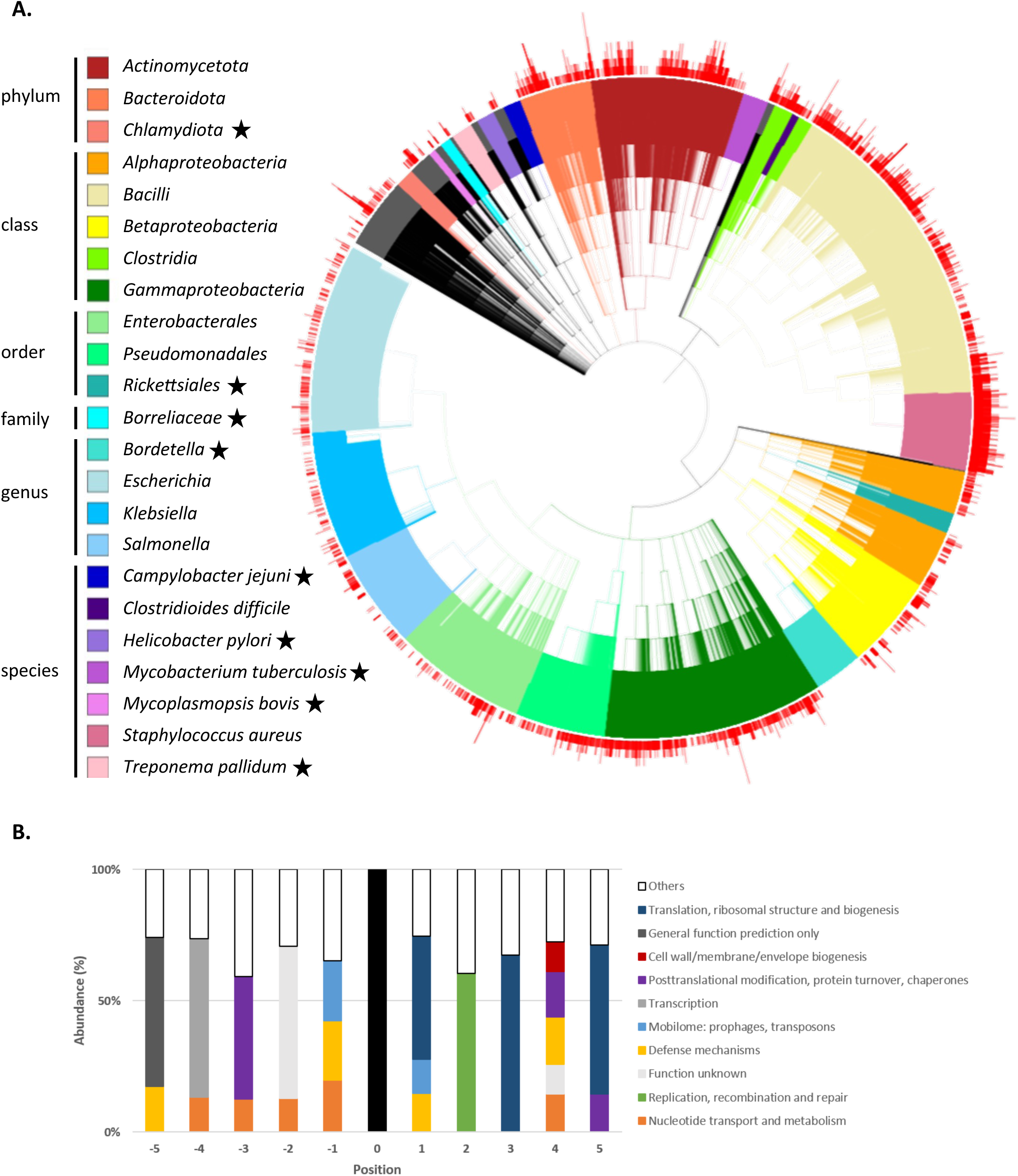
- Distribution and environment of AbiF/Abi_2 systems in bacteria. (**A**) Distribution of AbiF/Abi_2 systems in 46,864 bacterial genomes from different phyla. The phylogenetic tree was made with iTOL v7 server by downloading the 3 files: the phylogenetic tree in newick format, the count of AbiF systems per genome added to iTOL template_simplebar, and the colors added to iTOL template_color_strip (supplementary data). Colors were defined according to taxon levels to highlight *C. difficile* and *S. aureus* and visualize taxon with no AbiF/Abi_2 system. Other taxonomic levels were colored when they constituted a balanced group according to the number of genomes in which the AbiF-like system was searched, and those remaining are shown in black. Red bars correspond to the number (1 up to 8 copies) of *abiF*/*abi_2* genes in genomes. Black stars correspond to bacteria order/family/species without *abiF*/*abi_2* genes. (**B**) Environment of *abiF/abi_2* genes in bacterial genome. Plot of genes located at gene positions −5 to −1 (upstream) and +1 to +5 (downstream) of the *abiF/abi_2* (position 0). Genes that are the most frequently located at each position are represented by the colored portions, where color identifies categories of biological function. Genes present with an abundance of less than 10% were grouped in the “others” category.

### Identification of a ncRNA associated with the putative AbiF_Cd_ system

Comparative analysis of RNA-seq data from the *C. difficile* strain R20291 (ribotype 027) and the reference strain 630Δ*erm* grown to the late exponential phase in TY medium (7) revealed the presence of a potential ncRNA gene specific to the R20291 strain and located upstream of the *abiF_Cd_* gene. In particular, RNA-seq reads were detected in the intergenic region between the *holin* and *abiF_Cd_* genes of the R20291 strain (Fig. 3A). In agreement with RNA-seq data, Northern blot analyses on RNA extracts from the R20291 strain revealed the presence of a major transcript of about 100 nt, that we named RCd22, as well as a less abundant and longer transcript (Fig. 3B). The profiling of RCd22 expression during growth showed that the maximum level of expression is reached during late exponential phase (6h of growth in TY medium) (Fig. 3B). The laboratory reference strain 630Δ*erm* lacks the phi027 prophage and as expected, no RNA could be detected by Northern blot using a RCd22-specific probe (Fig. 3B). *In silico* analysis of the corresponding region in the R20291 genome revealed the presence of a Sigma A-dependent promoter consensus upstream of the RCd22 sequence. A predicted Rho-independent terminator was also detected in the RCd22 3’ end, in concordance with the estimated size of about 100-130 nucleotides for the detected transcripts (Fig. 3C). The RCd22 gene transcriptional start site (TSS) was located 168 nucleotides upstream of the *abiF_Cd_* ATG start codon associated with a ribosome binding site (AGGTGA) for initiation of translation. Moreover, no promoter could be predicted between the RCd22 terminator and the *abiF_Cd_* gene, suggesting that RCd22 and *abiF_Cd_* are co-transcribed. To map the 5’ and 3’ extremities of transcripts derived from this region, we performed a 5’/3’ Rapid amplification of cDNA ends (5’/3’ RACE) experiment. The major 5’-end was mapped to the A nucleotide position associated with the consensus elements of SigA-dependent promoter. Several 3’ extremities were mapped just upstream or inside the terminator hairpin structure followed by the four-U stretch that corresponds to the full-length 132 nt-transcript and shorter degradation or processing products ranging from 88 to 115 nt in length (Fig. 3C). The major 3’-end was located 8 nt upstream of the terminator stem-loop corresponding to the 98-nt transcript. This approach allowed us to confirm the presence of two groups of transcripts sharing the same TSS, a short one ranging from 88 nt to 115 nt in length and corresponding to the RCd22 sequence and a long one of 1,086 nt corresponding to the transcriptional readthrough leading to co-transcription of RCd22 and *abiF_Cd_* genes (Fig. 3C). RNA-seq data showed that the short form including the RCd22 sequence alone and lacking the *abiF_Cd_* coding sequence is mainly transcribed under laboratory conditions (Fig. 3A). These data suggest that the ncRNA RCd22 could regulate the expression of the downstream *abiF_Cd_* gene through a “riboswitch-like” mechanism associated with premature termination of transcription or maturation. Moreover, a conserved *abiF* motif (Rfam: RF03085) has been identified in RCd22. This motif is associated with putative *abiF* genes in various bacterial species and especially in environmental isolates of *C. difficile* (29). In RCd22, one copy of this motif is located between positions 9 and 41 according to the TSS, and a second copy is also present between positions 61 and 95 (Fig. 3D). A RCd22 sequence was also found upstream of the *abiF_Cd_* homologous genes in phages φC2 and φCD2301. This conserved co-localization of RCd22 and *abiF* suggests a functional link between RCd22 and the putative AbiF system in *C. difficile*.

**Fig. 3.**
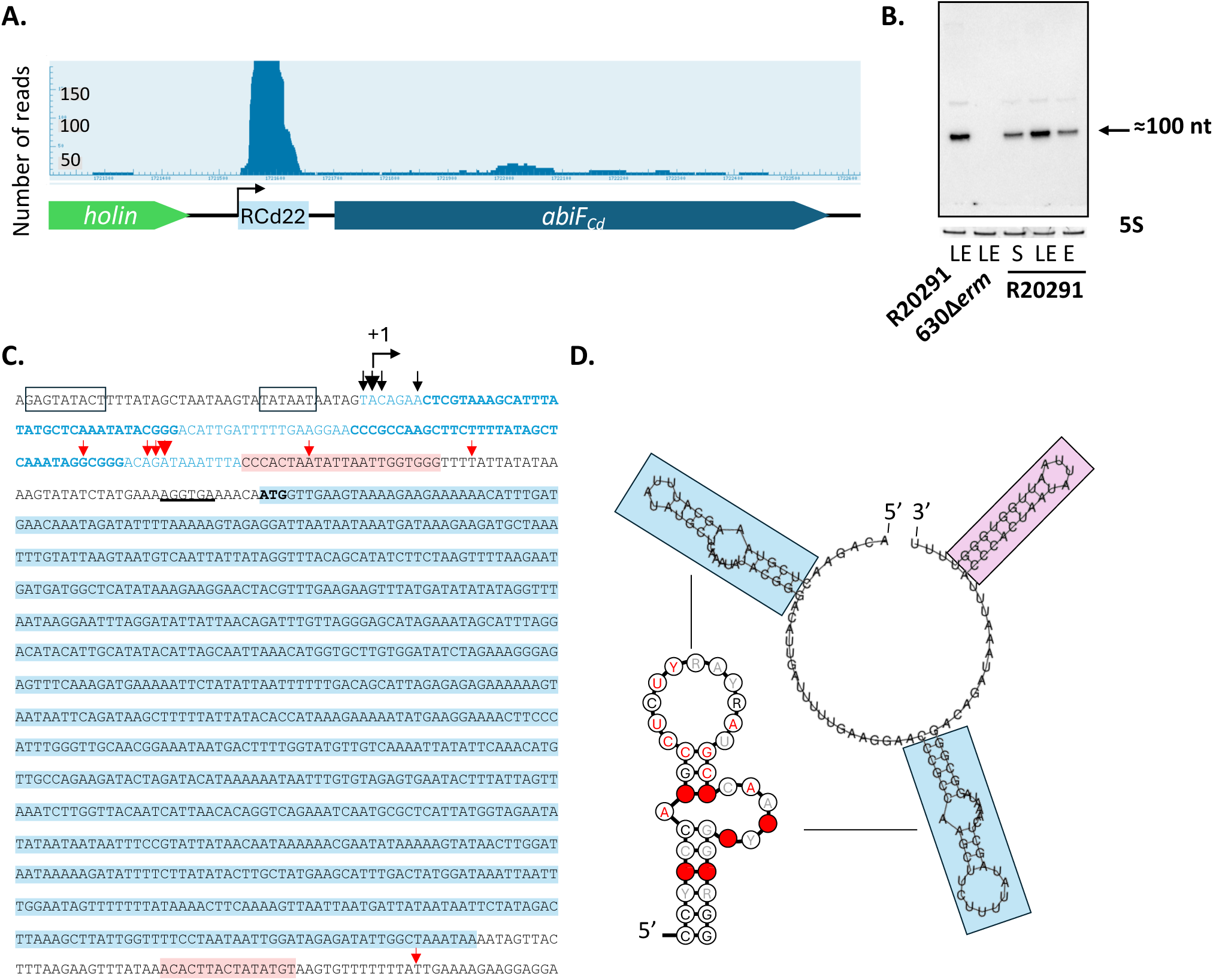
- A ncRNA RCd22 is associated with the hypothetical AbiF_Cd_ system. (**A**) RNA sequencing of *C. difficile* R20291 strain showing transcript reads of a ncRNA, named RCd22, upstream of the *abiF_Cd_* gene, encoding a hypothetical AbiF_Cd_ protein. (**B**) Detection of RCd22 by Northern Blot in the R20291 strain and the absence of the signal in the *C. difficile* reference laboratory strain 630Δ*erm*. RNA samples were extracted from the R20291 strain grown at exponential phase (E, 4h of growth), late exponential phase (LE, 6h of growth) or entry into stationary phase (S, 10h of growth) and from the 630Δ*erm* strain grown at late exponential phase (LE, 6h of growth). 5S rRNA is used as a loading control. (**C**) Detection of two groups of transcripts by 5’/3’RACE experiments in the R20291 strain. Short transcripts corresponding to RCd22 and a long transcript corresponding to the co-transcription of RCd22 and *abiF_Cd_.* Light blue letters correspond to RCd22 and light blue bold letters correspond to the two *abiF* motifs; nucleotides shaded in light blue correspond to the *abiF_Cd_* encoding gene; black boxes correspond to the consensus −10 and −35 elements of the predicted SigA-dependent promoter; nucleotides shaded in pink correspond to the predicted Rho-independent terminators; underlined letters correspond to the predicted ribosome binding site; black arrows depict 5’ ends and red arrows depict 3’ ends, the arrow size is proportional to the number of extremities identified by 5’/3’RACE. (**D**) Prediction of RCd22 secondary structure by RNAfold, and identification of two *abiF* conserved motifs (Rfam RF03085). Blue boxes correspond to *abiF* conserved motifs; pink box corresponds to the predicted Rho-independent terminator; R=A or G; Y=U or C; in *abiF* motif: red characters correspond to 97% nucleotide identity, black characters to 90% nucleotide identity and grey characters to 75% nucleotide identity, red circles correspond to 97% nucleotide conservation.

### Overexpression of *abiF_Cd_* induces growth defects in *C. difficile* and *E. coli* strains

The toxins from Abi systems generally induce bacterial death or growth arrest when overexpressed from plasmids (17,30,31). To determine whether the putative AbiF_Cd_ protein could be toxic for the cell, we analyzed the effect of its overexpression on the growth of *C. difficile* in broth and on agar. We first generated a plasmid for inducible overexpression of the *abiF_Cd_* gene under the control of the anhydrotetracycline (ATc)-inducible P_tet_ promoter (p/*abiF_Cd_*) (Fig. 4A). The construct was transferred by conjugation into the *C. difficile* strain 630Δ*erm* in which both φCD630-1 and φCD630-2 prophages were deleted to avoid potential interference (CD156). CD156 is used in this study as a control strain because it lacks the AbiF_Cd_ system. As shown in Figure 4B, after 4h of induction with the ATc inducer in liquid culture, bacterial growth was severely reduced for the strain carrying p/*abiF_Cd_* plasmid as compared to the strain carrying an empty vector. A similar growth defect was observed in the R20291 strain overexpressing *abiF_Cd_* from a plasmid, and also carrying the AbiF_Cd_ system on the phi027 prophage (Fig. S2A). The bacteriostatic effect of *abiF_Cd_* overexpression was further confirmed by monitoring CFU counts over time (Fig. S2B). No growth difference was observed for *C. difficile* strain CD156 carrying p/*abiF_Cd_* or an empty vector in the absence of ATc both in liquid medium (Fig. S3) and on TY plates (Fig. 4C). By contrast, the overexpression of *abiF_Cd_* caused a growth defect and reduced colony size on plates of the *C. difficile* strain carrying p/*abiF_Cd_* in the presence of ATc as compared to the control strain (Fig. 4C). To assess the toxic activity of AbiF_Cd_ in a heterologous host, we also analyzed the effect of its overexpression in *E. coli* grown in liquid culture. A severe growth defect was observed in the *E. coli* strain carrying the p/*abiF_Cd_* plasmid in the presence of ATc as compared to the control strain with empty vector (Fig. S2C). Altogether, these results demonstrate the toxic activity of the AbiF_Cd_ protein that induces growth reduction without cell death in *E. coli* and *C. difficile*.

**Fig. 4.**
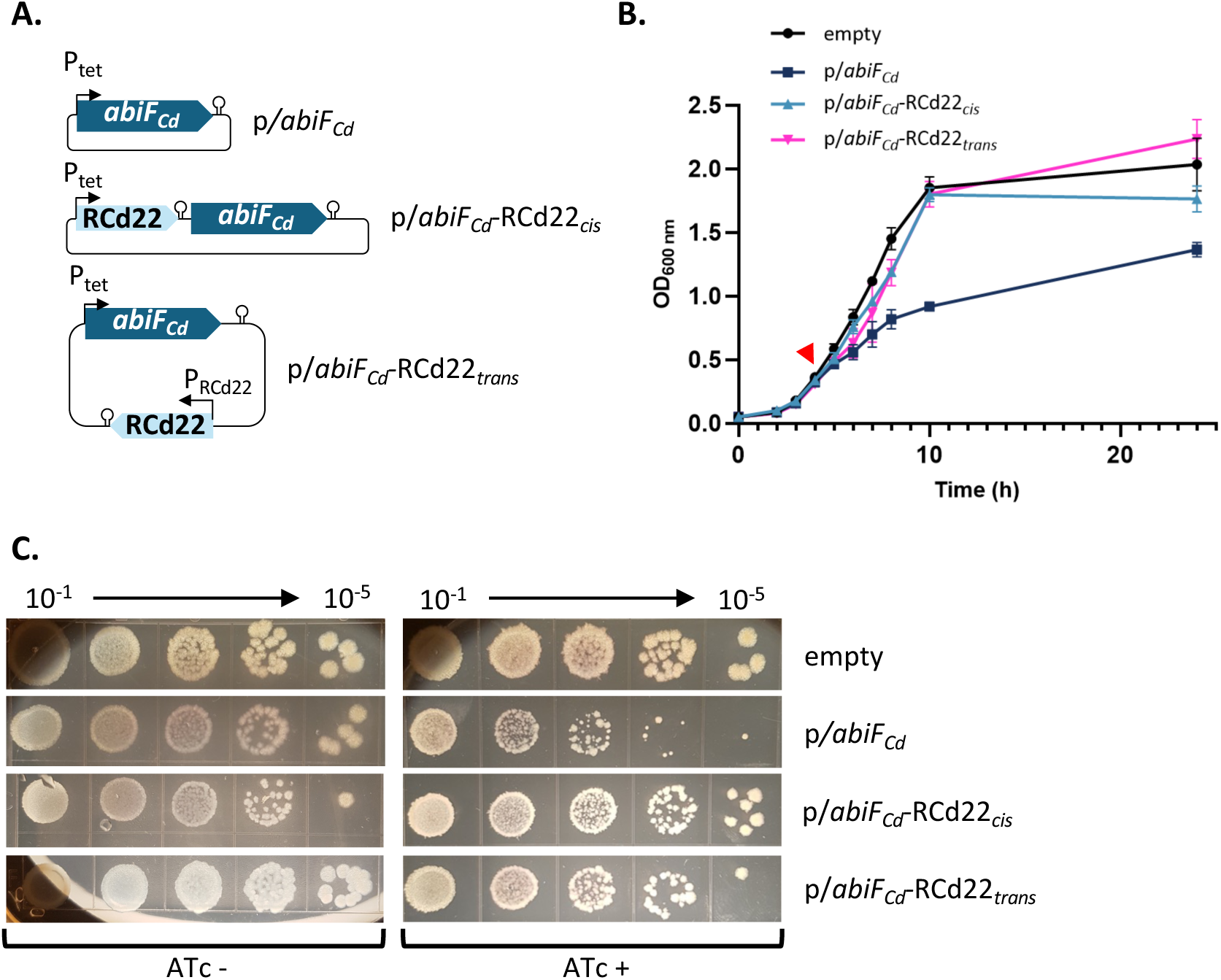
- Impact of *abiF_Cd_* gene overexpression on *C. difficile* growth. (**A**) Schematic representation of plasmids used for these experiments. p*/abiF_Cd_* corresponds to pDIA6103 plasmid carrying *abiF_Cd_* under the control of the inducible P_tet_ promoter; p/*abiF_Cd_-*RCd22*_cis_* carries both RCd22 and *abiF_Cd_* under the control of P_tet_ promoter, separated by a Rho-independent terminator; p/*abiF_Cd_-*RCd22*_trans_* carries *abiF_Cd_* under the control of P_tet_ promoter and RCd22 under the control of its native promoter (P_RCd22_). (**B**) Growth curve of *C. difficile* strain CD156 (630 Δ*erm* ΔphiCD630-1ΔphiCD630-2) carrying pDIA6103 empty plasmid (empty), p*/abiF_Cd_,* p/*abiF_Cd_-*RCd22*_cis_* or p/*abiF_Cd_-*RCd22*_trans_*. Induction of the P_tet_ promoter by 250ng/mL ATc is indicated by the red arrow. Plotted values represent means and error bars represent standard error of the means (N = 3 biologically independent samples). (**C**) Spot assay of *C. difficile* CD156 strains carrying the different plasmid constructions on TY agar plates supplemented with Tm and with or without 250ng/mL ATc inducer. Pictures are taken after 48h incubation at 37°C.

### RCd22 down-regulates *abiF_Cd_* expression at the transcriptional level and represses AbiF_Cd_ toxic activity

To investigate the regulation mechanism of the putative AbiF_Cd_ system in the R20291 strain, we first assessed the possibility of a transcriptional control. We deleted the RCd22 sequence from the R20291 genome, including both *abiF* motifs and the predicted terminator, but kept the promoter of RCd22 and the RBS of *abiF_Cd_*. The absence of RCd22 expression in the ΔRCd22 strain was confirmed by qRT-PCR (Fig. S4A). We then measured the level of *abiF_Cd_* expression by qRT-PCR in the absence or the presence of RCd22 (Fig. 5A). In the strain deleted for RCd22, we observed a 17-fold increase in *abiF_Cd_* expression as compared to the wild-type control strain. This shows down-regulation of *abiF_Cd_* expression by RCd22. Accordingly, the deletion of RCd22 also induced a growth defect and a decreased growth yield in the R20291 strain (Fig. S4B). The stronger growth defect observed in the 630Δ*erm* strain correlated with a higher level of *abiF_Cd_* expression from the plasmid (Fig. 4B, Fig. S5A). To further analyze the effect of RCd22 on *abiF_Cd_* expression, the RCd22 sequence was fused to the *phoZ* reporter gene on a plasmid. The expression was under the control of a constitutive promoter (P_cwp2_) and the alkaline phosphatase activity was measured (Fig. 5B). The plasmid carrying the RCd22-*phoZ* transcriptional fusion (p/RCd22-*phoZ*) and the control plasmid with *phoZ* under the control of the P_cwp2_ constitutive promoter (p/*phoZ*) were conjugated in four different strains: the 630Δ*erm* naturally lacking the AbiF system, the R20291 wild-type strain carrying the phi027 prophage and the AbiF system, the R20291 Δphi027 strain deleted for the prophage, and the R20291 ΔRCd22 strain deleted from the RCd22 gene. The presence of RCd22 upstream of the *phoZ* gene severely reduced the alkaline phosphatase activity in all strains analyzed (Fig. 5B). These results suggest that RCd22 down-regulates *abiF_Cd_* expression at the transcriptional level independently from the genetic background used. They also suggest that the presence of the Rho-independent terminator at the 3’-end of RCd22 leads to a strong termination of transcription, thereby reducing the read-through transcription of the downstream gene *abiF_Cd_*.

**Fig. 5.**
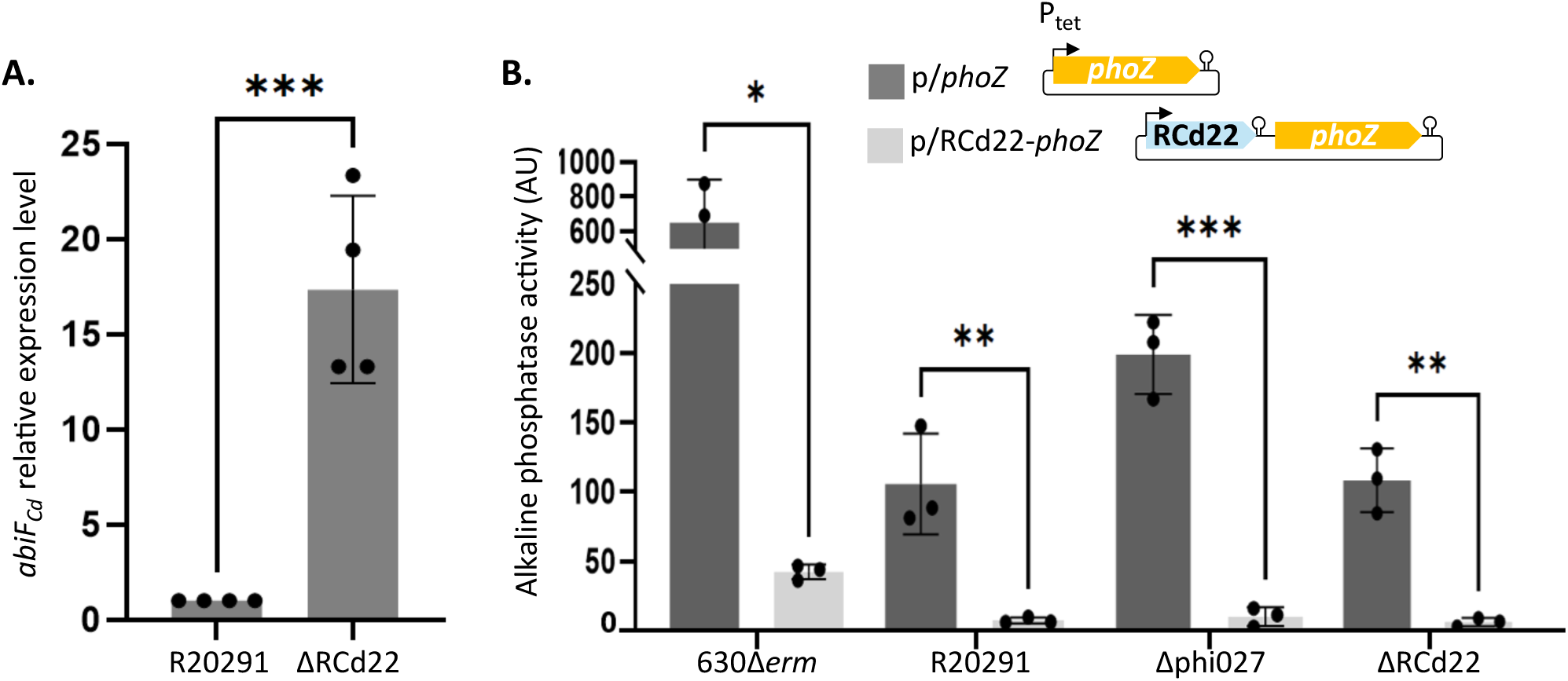
- RCd22 down-regulates *abiF_Cd_* gene expression at the transcriptional level. (**A**) *abiF_Cd_* expression level measured by qRT-PCR, in R20291 ΔRCd22 compared to WT (N=4, biologically independent samples). (**B**) Alkaline phosphatase activity of the RCd22-*phoZ* reporter fusion (p/RCd22-*phoZ*) under the control of the constitutive P_cwp2_ promoter, compared to empty plasmid (p/*phoZ*) in different genetic backgrounds of *C. difficile*. Values represent the mean standard error of the mean (N = 3 biologically independent samples). Asterisks indicates significant differences (*t* test): * indicates a p-value<0.05; ** indicates a p-value<0.01; *** indicates a p-value<0.001.

We then tested the role of RCd22 as a repressor of AbiF_Cd_ toxic activity. The RCd22 sequence was cloned in the p/*abiF_Cd_* plasmid, upstream of the *abiF_Cd_* gene to give p/*abiF_Cd_*-RCd22*_cis_* (Fig. 4A). In agreement with the ability of RCd22 to down-regulate *abiF_Cd_* expression, the presence of RCd22 in *cis* repressed the toxic activity of AbiF, leading to a complete reversion of the growth defect in liquid culture in the presence of ATc inducer (Fig. 4B). Similar results were obtained on agar plates since the presence of RCd22 in *cis* led to the recovery of normal colony size and normal growth (Fig. 4C). To get insights into the RCd22 mode of action, we also cloned it under the control of its own promoter *in trans* in a different location on the plasmid (p/*abiF_Cd_*-RCd22*_trans_*). Co-expression of the RCd22 *in trans* along with the *abiF_Cd_*, also led to the inhibition of the toxic activity of AbiF_Cd_ (Fig. 4B, C). The transcript abundance of each gene was controlled by qRT-PCR after 4h of induction (Fig. S5). In agreement with reporter fusion assays (Fig. 5B), RCd22 expression *in cis* negatively impacted the expression of *abiF_Cd_*, while RCd22 expression *in trans* did not affect *abiF_Cd_* expression (Fig. S5). These results clearly show that RCd22 can act both *in cis* and *in trans* to control AbiF_Cd_ toxic activity.

### RCd22 interacts with the AbiF_Cd_ protein

We then investigated the mechanism of regulation *in trans* by RCd22 of *abiF_Cd_*. We wondered whether RCd22 interacts with *abiF_Cd_* mRNA or AbiF_Cd_ protein to counteract its toxic activity. To discriminate these two hypotheses and to identify all mRNA and protein targets of RCd22, we used an MS2-Affinity Purification approach coupled with RNA-Sequencing (MAPS) or Mass Spectrometry (32). For this experiment, we first added an MS2-tag to the 5’-end of the RCd22 ncRNA and integrated this construct into a plasmid under the control of an inducible P_tet_ promoter (Fig. 6A). We conjugated the resulting plasmid into the R20291 ΔRCd22 strain and induced the expression of the MS2-RCd22 ncRNA during exponential growth phase in the presence of ATc inducer. We then collected all mRNA and protein targets by affinity purification using an MBP-MS2 protein complex that interacts with the MS2-tag. The mRNA targets were identified by RNA-sequencing while protein targets were identified by mass spectrometry. We compared these results with the strain carrying the plasmid with the MS2-tag alone as a control, to eliminate targets interacting with MS2-tag. As expected, RCd22 appeared as the most enriched RNA in this analysis. In addition to CDR20291_RS16617 corresponding to a lysine riboswitch upstream of the *asd CDR20291_3084* gene, many potential mRNA targets were identified with this technique, including mRNA involved in nucleic acid metabolism (Fig. S6A, Supplementary Table S4). We then used IntaRNA to predict *in silico* the RNA-RNA interactions between RCd22 and its potential targets. We observed that single strand regions of RCd22 base-paired with the ORF of mRNA potential targets (Fig. S6B). Six mRNAs, including *CDR20291_1462* containing the predicted sites of interaction with RCd22, were *in vitro* transcribed and mixed with radiolabeled RCd22. For three of them (*CDR20291_1558, 2768* and *3357*), the transcribed RNA was divided into a 5’ and a 3’ part for interaction analysis. Gel shift assays did not reveal any formation of RNA-RNA complexes (Fig. S6C), except for the *CDR20291_1558* 3’ fragment, where a complex with RCd22 appeared, exhibiting a weak affinity (Fig. S6C). This indicates that RCd22 generally not act on mRNA targets *in trans*. For the protein targets, mass spectrometry analysis of MAPS protein fraction identified the AbiF_Cd_ protein as the main protein target of RCd22 (Fig. 6B, Supplementary Table S5), with an enrichment log2 fold change (FC) of 6. The enrichment score of other potential protein targets was under a log2(FC) of 2.53. This result indicates that RCd22 interacts with the AbiF_Cd_ protein. We concluded that the main target of RCd22 is the AbiF_Cd_ protein. Thus, the ncRNA could act as an antitoxin by directly targeting and neutralizing the AbiF_Cd_ protein, similar to type III TA systems.

**Fig. 6.**
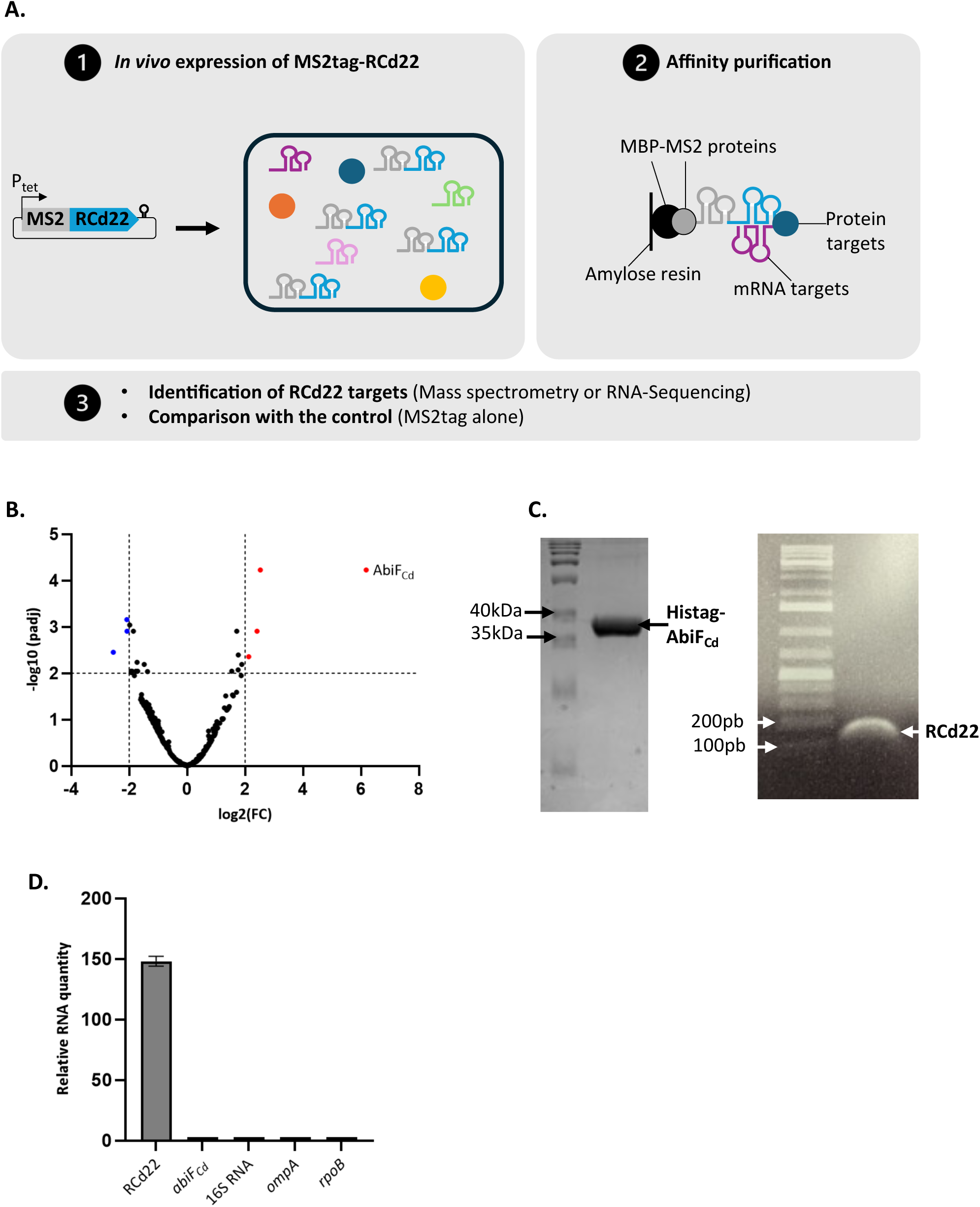
- RCd22 interacts with the AbiF_Cd_ protein. (**A**) Schematic representation of MS2 Affinity Purification experiment coupled with mass spectrometry or RNA sequencing. (**B**) Volcano plot of RCd22 protein targets. Red dots indicate enriched proteins targeted by RCd22 compared to the control. (**C**) Co-purification of RCd22 with Histag-AbiF_Cd_. SDS-PAGE of Histag-AbiF_Cd_ purified fraction from HisTrap affinity column and agarose gel after RNA extraction from Histag-AbiF_Cd_ purified fraction. (**D**) Representative result of at least two independent experiments for qRT-PCR of RNA extracted from Histag-AbiF_Cd_ purified fraction. Several targets were tested, relative RNA quantity is estimated according to the standard curve with this formula: 10^((Cq-intercept)/slope).

To investigate the RCd22-AbiF_Cd_ interaction *in vitro*, we first purified the 10xHis-tagged AbiF_Cd_ protein expressed in *E. coli* (Fig. S7) and confirmed that this recombinant protein conserved its toxic activity on cell growth (Fig. S7A). Since the AbiF_Cd_ protein is toxic for *E. coli* (Fig. S2C and Fig. S7A) and its production was not efficient (Fig. S7B), we co-expressed RCd22 with *abiF_Cd_* to counteract the toxic activity during *abiF_Cd_* induction and improve AbiF_Cd_ production (Fig. S7C). The analysis of elution fractions from the first HisTrap affinity purification step on SDS-PAGE revealed a protein of the expected size at 34.2kDa (Fig. 6C). In agreement with MAPS results, we were able to co-purify the ncRNA RCd22 by targeting AbiF_Cd_ in the absence of nuclease treatment. Indeed, RNA extraction followed by agarose gel and RT-qPCR analysis detected RCd22 in the HisTrap column elution fractions (Fig. S7D, Fig. 6C, D). However, despite numerous attempts, we were unable to get the purified AbiF_Cd_ protein exempt of RCd22, suggesting the formation of stable RNA-protein complex with low dissociation rate.

### The RCd22 secondary structure and both *abiF* motifs are important for neutralization of AbiF_Cd_ toxic activity

*In silico* predictions suggested that RCd22 is a highly structured RNA with two conserved *abiF* motifs folding into stable hairpin structures that could constitute important functional elements of the antitoxin RNA. To test this hypothesis, we first introduced mutations in both RCd22 *abiF* motifs and then tested their effects on the antitoxin activity of RCd22 (Fig. 7A). The first type of mutation (mut1, Fig. 7A) was designed to affect the stem structure stability in both RCd22 *abiF* motifs (*abiF1* and *abiF2*) by replacing a G by a C, thus removing one base pair within this motif. The second type of mutation (mut2, Fig. 7A) was designed to change the sequence without affecting the structure of RCd22. For this, two C and G nucleotides in the stem of the *abiF1* and *abiF2* motifs were inverted, thus keeping the base-pairing inside the motif. With these two types of mutations, we sought to get information about the dependence of the RCd22 structure or sequence for neutralization of the toxicity of AbiF_Cd_. These mutations were introduced into plasmids co-expressing *abiF_Cd_* and RCd22 *in cis* or *in trans.* The expression of both *abiF_Cd_* and RCd22 was validated by qRT-PCR analysis after their introduction into *C. difficile* by conjugation (Fig. S5). On the one hand, the RCd22 mut2 variants that retained the stem-loop structures down-regulated the expression *abiF_Cd_ in cis,* similar to WT RCd22 expressed *in cis*. On the other hand, the RCd22 mut1 variants, carrying destabilized stem structures, were no longer able to repress *abiF_Cd_* expression (Fig. S5). The capacity of mutated RCd22 to neutralize the toxic activity of AbiF_Cd_ was then tested in liquid medium and on agar plates. RCd22 with mutations destabilizing the stem structure (RCd22mut1) were no longer able to neutralize the toxicity of AbiF*_Cd_*. Indeed, neither the growth defect nor the colony size could be restored, in contrast to the normal growth of strains carrying unmodified RCd22 constructs (Fig. 7B, C). This effect was observed for both constructs expressing the RCd22 *in cis* and *in trans*. A similar action of RCd22 was expected for both constructs *in trans,* through the produced RNAs that interact with the toxin. However, for the *cis*-located RCd22 mutant variants, the introduced mutations could destabilize the stem structure and affect the conformation of the terminator, thus leading to *abiF_Cd_* transcription and AbiF_Cd_ translation. We thus concluded that the stem structure within the two RCd22 *abiF* motifs was important for the interaction with the AbiF_Cd_ protein and neutralization of its toxic activity. On the contrary, compensatory mutations that only changed the RCd22 sequence inside the stem without altering its structure (RCd22mut2) did not affect neutralization of the toxic activity *in cis* and *in trans* (Fig. 7B, C). These nucleotides might not be important for the interaction with AbiF_Cd_, as they are base-paired in the stem and are probably not accessible for the interaction with the protein. Rather, the structure of these two motifs could be important to keep nucleotides in between accessible for interaction with the protein. This result is consistent with the *in silico* prediction of the RCd22-AbiF_Cd_ complex (see below and Fig. 8).

**Fig. 7.**
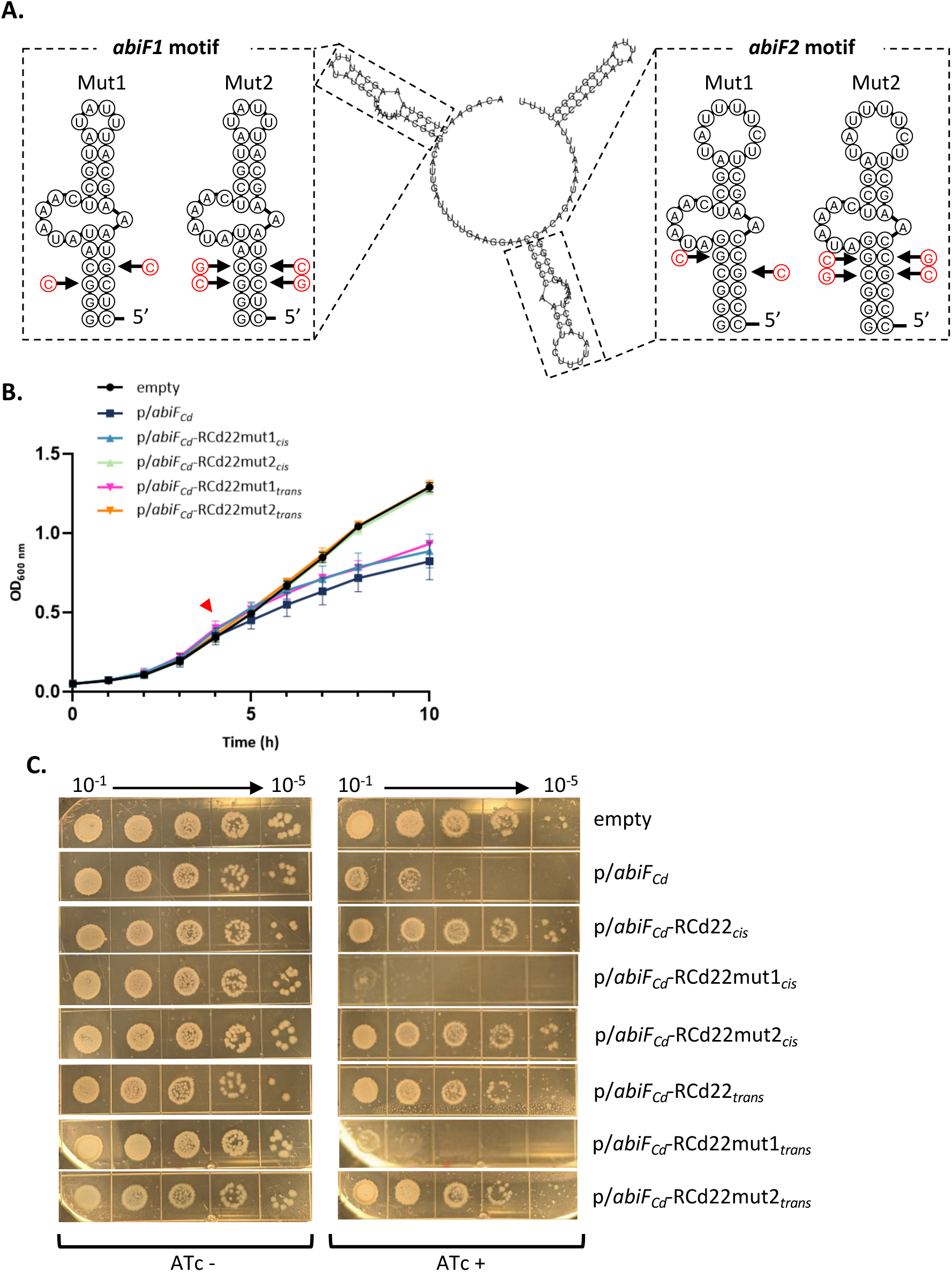
- Importance of *abiF* motifs and RCd22 structure for AbiF_Cd_-RCd22 interaction. (**A**) Schematic representations of mutations introduced in the RCd22 conserved *abiF* motifs. The predicted nucleotides potentially contributing to AbiF_Cd_-RCd22 interaction are indicated in red. (**B**) Growth curve of *C. difficile* strain CD156 carrying either the pDIA6103 empty plasmid (empty), or expressing *abiF_Cd_* under the control of the inducible P_tet_ promoter (p*/abiF_Cd_)* or co-expressing *abiF_Cd_* and RCd22 (*in cis* or *in trans,* with or without mutations 1 or 2). Induction of the P_tet_ promoter by 250ng/mL ATc is indicated by the red arrow. Plotted values represent the mean standard error of the mean (N = 3 biologically independent samples). (**C**) Spot assay of *C. difficile* CD156 strains on TY agar plates supplemented with Tm and 250ng/mL ATc inducer. Pictures are taken after 48h incubation at 37°C.

**Fig. 8.**
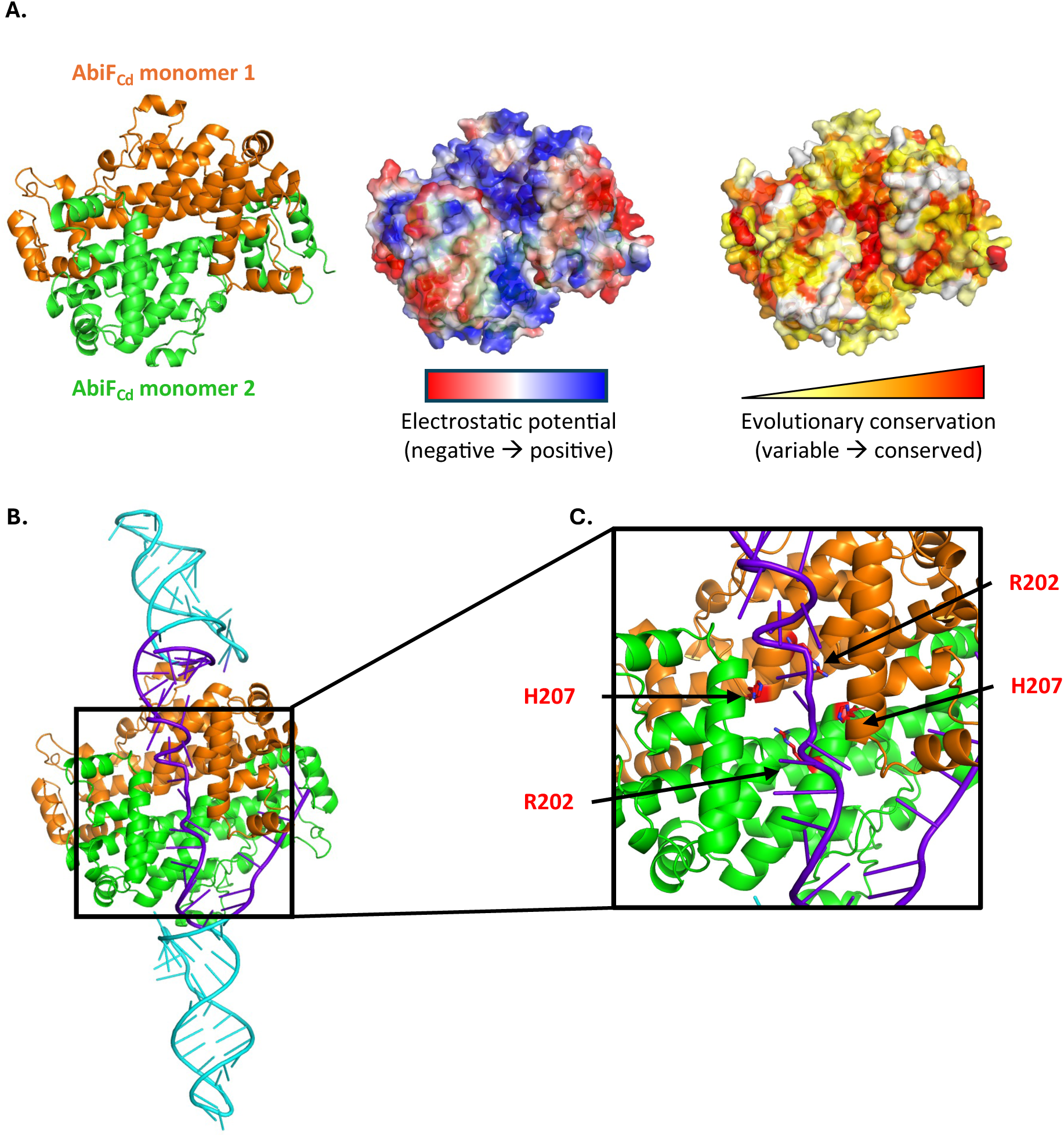
– *In silico* structural prediction of the AbiF_Cd_-RCd22 complex. (**A**) Prediction of the homodimer AbiF_Cd_ structure by AlphaFold2-Multimer. The AbiF_Cd_ homodimer complex is represented in orange and green (left). The electrostatic potential (from red to blue, negatively to positively charged) and evolutionary conservation (from white to red, most variable to most conserved) were mapped on the surface of the homodimer and visualized with Pymol. (**B**) Prediction of AbiF_Cd_-RCd22 interaction by AlphaFold3. The RCd22 sequence is in purple with the two *abiF* motifs in cyan. (**C**) Prediction of important amino acids, R202 and H207, for AbiF_Cd_ mutagenesis. Amino acids of both AbiF_Cd_ monomers are represented as sticks and indicated in red.

### *In silico* prediction of RCd22-AbiF_Cd_ complex structure and identification of key residues for toxicity

We further analyzed the RCd22-AbiF_Cd_ interaction to determine which amino acids and nucleotides are involved in this complex formation. We predicted the AbiF_Cd_ protein structure using AlphaFold2-Multimer (Fig. 8A) and the structure of the RCd22-AbiF_Cd_ complex using AlphaFold3 (Fig. 8B). The AlphaFold2 structural model for the AbiF_Cd_ protein (best out of 5 models, see Methods) suggested with very high confidence that it could form a homodimer (predicted TM-score (pTM) = 0.946 and interface predicted TM-score (ipTM) = 0.935). The AlphaFold3 structural model (best out of 25 models, see Methods) including two copies of AbiF_Cd_ and one copy of a short (98-nt) abundant transcript of RCd22 (Fig. 8B) displayed overall good confidence (pTM = 0.8 and ipTM = 0.71). This was mostly driven by the very high confidence of the protein homodimer prediction (protein chain pTM = 0.87 and protein-protein ipTM = 0.87). The protein-protein homodimer predicted by AlphaFold3 could be superimposed almost exactly on the AlphaFold2 prediction (root mean square deviation (RMSD) = 0.5 Å). The RNA molecule in the AlphaFold3 prediction had much lower confidence than the proteins (RNA pTM = 0.12 and RNA-protein ipTM = 0.17-0.18). This is not uncommon in our experience with AlphaFold3 predictions involving RNA molecules with no close homologs in the Protein Data Bank. Our confidence in the RNA and protein-RNA predictions was nevertheless reinforced by several observations: (i) the secondary structure of the RNA stem loops in the AlphaFold3 model matched exactly the secondary structure predicted by RNAfold (Fig. 3D); (ii) the two *abiF* conserved motifs interacted with the same zones of each monomer, although the exact binding modes were different (which was expected given the sequence difference between the two motifs); and (iii) the interaction surface created by the two AbiF_Cd_ monomers presented a conserved, positively charged groove highlighted in the electrostatic mapping (Fig. 8A), where the single strand RNA region in-between the two *abiF* motifs is lodged in the model.

The above-mentioned conserved, positively charged groove is an interesting candidate for interaction with nucleic acids and RNase activity of toxin AbiF_Cd_ (Fig. 8). To validate the importance of this region for AbiF_Cd_ toxicity, we defined two conserved arginine R202 and histidine H207 residues (Fig. 8C) within the basic groove formed at the AbiF_Cd_ dimer interface for targeted mutagenesis (Fig. 8C). We introduced amino acid substitutions to replace R202 and H207 residues either by negatively charged aspartic acid or by neutral alanine. The corresponding plasmids were used to overexpress these protein variants and test their impact on *C. difficile* growth (*abiF_Cd_*mut1 and *abiF_Cd_*mut2 in Fig. 9). In the presence of ATc inducer, both types of mutations led to the loss of toxic activity. Little to no growth defect was observed in liquid and solid culture as compared to the severe growth inhibition following overexpression of the native AbiF_Cd_ protein (Fig. 9).

**Fig. 9.**
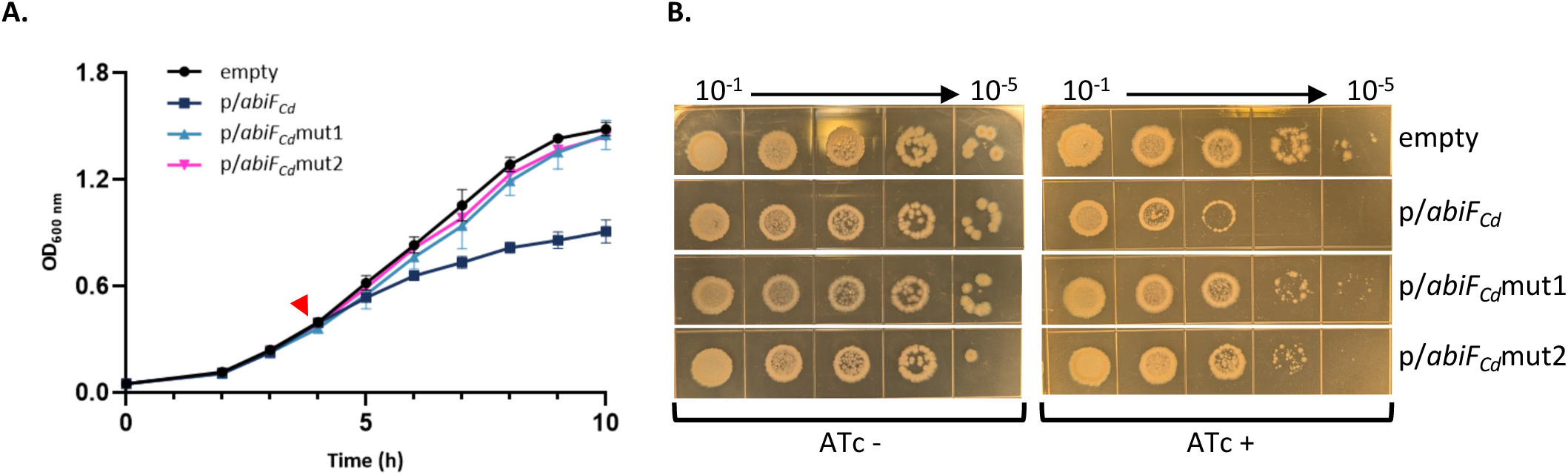
– Importance of R202 and H207 residues for AbiF_Cd_ toxic activity. (**A**) Growth curve of *C. difficile* strain CD156 carrying either the pDIA6103 empty plasmid (empty), or expressing *abiF_Cd_* mutated variants (substitutions R202D+H207D, named *abiF_Cd_*mut1 or R202A+H207A, named *abiF_Cd_*mut2) under the control of the inducible P_tet_ promoter. Induction of the P_tet_ promoter by 250ng/mL ATc is indicated by the red arrow. Plotted values represent the mean standard error of the mean (N = 3 biologically independent samples). (**B**) Spot assay of *C. difficile* CD156 strains on TY agar plates supplemented with Tm and 250ng/mL ATc inducer.

## Discussion

In this study, we provide the first characterization of an AbiF-like system in Clostridia and unravel its regulation mechanism by the ncRNA RCd22. Abi systems are generally defined as bacterial defense mechanisms against phages, where bacteria die or stay in a dormancy state before the infecting phage can complete its lytic cycle, thus protecting the bacterial population (1). Intriguingly, AbiF_Cd_ is encoded within the well-conserved phi027 prophage in the R027 hypervirulent strains of *C. difficile* and in two other *C. difficile* prophages, φC2 and φCD2301. AbiF_Cd_ belongs to the Abi_2 superfamily and AbiD/F group, first identified and characterized in *L. lactis* pNP40 plasmid (12). AbiD, AbiD1 and AbiF systems superimpose structurally, giving a single AbiD/F group (13) involved in phage resistance (12,33). This protein family has structural similarities to the HEPN (Higher Eukaryotes and Prokaryotes Nucleotide-binding) domains of RNA-guided RNase Cas13 in type VI RNA-targeting CRISPR-Cas systems (34). HEPN RNases are widespread in prokaryotic defense systems including type II and type VI CRISPR-Cas, as well as type II and type VII TA systems and are parts of the eukaryotic RNA processing and degradation machineries (35,36). It was suggested that Cas13 nuclease domain evolved from ancestral AbiD/F ribonuclease with potential RNA-guided capabilities (34) that could be related to a conserved *abiF* RNA motif of 36-37 nt associated with AbiF system (RNA family: RF03085) (29). During the preparation of this manuscript, in agreement with the proposed model, Zilberzwige-Tal *et al.* reported biochemical and structural characterization of the evolutionary origins of CRISPR-Cas13 from AbiF through a miniature Cas13e (37). They experimentally solved the structure of *Prevotellaceae bacterium* AbiF (PbAbiF), which forms a homodimer, in complex with two copies of a ncRNA. Comparison of this experimental structure with our AlphaFold3 model of the RCd22-AbiF_Cd_ complex (Fig. S8) shows very good superimposition of the protein homodimer (RMSD = 3.1 Å). The interaction surfaces of the ncRNA on PbAbiF are very similar to the interaction surfaces of the *abiF* motifs of RCd22 on AbiF_Cd_, despite some divergence in the ncRNA structures. The presence of two *abiF* motifs in a single RCd22 molecule means that the single-strand RNA region in between the two motifs has no equivalent in the PbAbiF/ncRNA structure. The amino acids R202 and H207 in AbiF_Cd_ are structurally aligned with R210 and H215 in PbAbiF, and the R210A+H215A variant of PbAbiF was shown in (37) to be inactive.

A Blast search for *abi_2*/*abiF* domains in bacterial genomes showed that Abi_2/AbiF domain is widely distributed with up to eight *abi_2*/*abiF* genes per genome. Surprisingly, some of the putative *abi_2/abiF* genes were located within prophages, downstream from the lysis module composed of a holin-endolysin operon, whereas it is located inside the lysis module in *C. difficile.* This location could be linked with the function or activity of AbiF_Cd_ system in *C. difficile*. Most Abi systems described in the literature are plasmid-encoded (38), but some were found within prophages. For example, two Abi systems were reported in *L. lactis* prophages with ORF852 encoding a putative AbiF protein downstream from the endolysin gene of the t712 prophage and ORF2248 encoding an AbiLi-like protein as a part of a two-component abortive infection system within *L. lactis* MG-4 prophage (39). Other anti-phage systems considered as Abi were described in *S. aureus* (26) and in *Mycobacterium* sp. prophages (27,40). Abi systems inside prophage genomes could function to protect from infection by other lytic phages or to stabilize the prophage in the bacterial chromosome, as demonstrated for type I toxin-antoxin systems (23,41,42).

The activity of AbiD/D1/F systems was first characterized in *L. lactis* plasmids as abortive phage infection modules also contributing in some cases to the low temperature stress response (12,15,16,33). Here we demonstrate that AbiF_Cd_ has a toxic activity in *C. difficile* and *E. coli*. We showed that overexpression of *abiF_Cd_* induced a bacterial growth defect in liquid medium and reduced colony size on plates, suggesting a bacteriostatic effect. Even though most Abi systems induce bacterial death, growth arrest was also observed as a strategy to let time for the bacteria to develop efficient defense against phage infection (31). Interestingly, in relation to the evolutionary origins of CRISPR-Cas13 from AbiF system, the Cas13 protein has a nonspecific RNase activity triggered by target recognition, and induces bacterial dormancy during phage infection, reminiscent to the abortive infection mechanism (43–45). This strategy seems to provide robust defense against phage infection but also prevents the emergence of CRISPR-resistant mutants (43). The collateral RNA degradation by Cas13a targets the anticodons in a subset of tRNAs, leading to the inhibition of protein synthesis and thus providing anti-phage defense (45). In addition, this tRNA cleavage indirectly triggers the activation of RNases from type II TA targeting mRNAs, contributing to the dormancy state (45). The conserved RNase motif RφxxxH, where φ is the polar residue N, H, or D, is associated with the catalytic site of HEPN RNases (35). Using AlphaFold protein modeling, we defined the amino acid substitutions targeting the conserved R202 and H207 residues within the RNase motif of AbiF_Cd_ of the basic groove formed at the AbiF_Cd_ dimer interface. When overexpressed on a plasmid in *C. difficile,* the AbiF_Cd_ protein variants mutated at these key positions lost their toxic activity, in agreement with the presence of the HEPN RNase **R**NQCA**H** motif. Further studies will be required to define the specificity of the RNase activity of AbiF_Cd_ contributing to its toxicity in *C. difficile* and in the heterologous *E. coli* host. The anti-phage activity of AbiF_Cd_ remains challenging to demonstrate in *C. difficile*. Indeed, all known *C. difficile* phages are temperate and lysogeny is frequent, therefore leading to significant lysogeny-mediated phage resistance. Based on its prophage localization, AbiF_Cd_ could also contribute to prophage maintenance as a common strategy associated with TA modules inside mobile genetic elements (46).

Since the AbiF_Cd_ is toxic for *C. difficile*, it must be tightly regulated under normal growth conditions. In this paper, we describe the RNA-based regulation mechanism of an AbiF-like system in *C. difficile* adding further evidence for the role of RNAs in the crossroads of phage-bacteria interactions. RNAs have emerged as crucial components of the numerous anti-phage defense strategies including CRISPR-Cas, TA and reverse-transcriptase-associated systems (47,48). The presence of a ncRNA, named RCd22, upstream of the *abiF_Cd_* gene was discovered by RNA-sequencing of the R20291 strain and further detected by Northern Blot. 5’/3’-RACE experiments suggest that RCd22 and *abiF_Cd_* genes are co-transcribed from the same SigA-dependent promoter. However, most of the transcription stops at a Rho-independent terminator upstream of the *abiF_Cd_* coding sequence, leading to the accumulation of a major and abundant transcript, RCd22. The presence of RCd22 within the 5’-untranslated-region of the *abiF_Cd_* mRNA impacts the expression of the downstream gene through premature transcription termination, similar to riboswitch-dependent regulations. We showed that this termination of transcription is strong enough to counteract the toxic effect of AbiF_Cd_ on bacterial cell. In addition to this antitoxin activity *in cis*, RCd22 was also able to counteract the AbiF_Cd_ toxic activity *in trans*. For the first time, we adapted the MAPS approach in *C. difficile* for RCd22 interactomics analysis and defined the direct RNA-protein interaction behind this antitoxin RCd22 activity *in trans.* We successfully identified the AbiF_Cd_ protein as the most enriched target of the ncRNA antitoxin. This powerful approach has been previously adapted in Gram-positive bacteria, for example to identify RNA partners of ncRNAs in *S. aureus* (49–51). By co-expressing the His-tag-AbiF_Cd_ protein with RCd22 in *E. coli*, we were also able to co-purify the ncRNA RCd22 with His-tag-AbiF*_Cd_*, providing additional experimental evidence for the RNA-protein complex formation. All our attempts to purify AbiF_Cd_ alone by removing the ncRNA from the complex were unsuccessful, suggesting a tight interaction between RCd22 and AbiF*_Cd_*, as previously reported for ToxIN type III TA RNA-protein complexes (52,53).

A similar two-step regulation mechanism for antitoxin action is operating in type III TA systems. In these TA systems, the antitoxin is a ncRNA transcribed from an array of short tandem repeats followed by the toxin protein coding gene (17,18). The antitoxin and toxin are co-transcribed and separated by a Rho-independent terminator, controlling the ratio of antitoxin RNA over the toxin mRNA. This balance between toxin mRNA and antitoxin ncRNA allows to keep the level of antitoxin in excess compared to the toxin, as the antitoxin is less stable than the toxin protein. The toxin protein activity is inhibited by the *trans*-acting antitoxin through specific RNA-protein interactions (17,18). However, the structure of both toxin and antitoxin from previously characterized type III TA systems differs from AbiF_Cd_ and RCd22 structural model. Indeed, in the most characterized ToxIN system in the plant pathogen *Pectobacterium atroseptica* (17,54) and in *E. coli* (55), protein-RNA interactions lead to a heterohexameric triangular assembly of three ToxN proteins with the interspersed pseudoknots of 36-nt ToxI RNAs (18). All other identified type III TA systems, i.e. AbiQ, TenpNI and CptNI, are homologous to ToxIN (56) with a structure similar to that of the AbiQ system (57), but with a completely different heterotetrameric quaternary organization for the CptNI complex due to extended antitoxin RNA size (58). Similar to previously characterized type III TA systems (17,18), the *abiF_Cd_* gene is preceded by an array of two repeated *abiF* RNA motifs within the RCd22 ncRNA gene. Antitoxin repeats have been defined as a key feature of type III TA systems (59). The ToxI or antiQ antitoxins from the ToxIN family are composed on an RNA motif of 34-36 nucleotides repeated from 2.8 to 5.5 times. The TenpNI and CptNI type III TA are associated with a longer RNA antitoxin of 50 and 45 nt repeated 2.1 times (59). At least one complete RNA repeat was essential for antitoxin activity *in vitro* for ToxIN (17) but the number of repeats required to keep the functionality varies between systems (60,61). In the ToxIN system, the repeated sequences in the transcript are cleaved by ToxN into individual 36-nt units followed by a self-assembly to 3 ToxN:3 ToxI complex (18). In contrast with other type III TA with toxin-mediated processing of antitoxin RNA, all the transcripts detected by RNA-seq, Northern blot and 5’/3’RACE analysis in *C. difficile* carried two *abiF* motifs of 33 and 35-nt in length interspaced by 19 nt. Potential processing sites were identified only upstream of the terminator stem-loop structure, with no cleavage between the *abiF* motifs. With respect to the evolutionary history of the AbiF-like systems at the origin of CRISPR-Cas13, such repeated stem-loop motif array inside the RCd22 antitoxin could mimic the CRISPR array organization, the duplication of *abiF* motif representing an intermediate step towards larger arrays of repeated motifs.

The *in silico* predictions suggest that two conserved *abiF* motifs of RCd22 (29) provide a stable structure that constitutes an important element for the functionality of the RCd22 antitoxin. We demonstrated that mutations affecting the structure of RCd22 led to an inactive form of antitoxin RNA unable to neutralize the toxic activity of AbiF_Cd_ *in cis* and *in trans*, while compensatory mutations had no impact on antitoxin activity. The overall RCd22 structure stabilized by two *abiF* motifs linked together could be important to keep the conformation of the RNA region in between accessible for the interaction with AbiF_Cd_ protein.

In conclusion, this work presents the identification of a new member of under-studied TA modules of type III operating through tight interaction between antitoxin RNA and toxic protein encoded inside the conserved prophage of the important human enteropathogen *C. difficile*. The association with the phage lysis module in several Gram-positive pathogens like *S. aureus, Streptococcus agalactiae* and *Listeria monocytogenes* is intriguing. Furthermore, the unique localization inside this module in *C. difficile* prophages deserves further investigations to better understand the role of AbiF system during interactions with phages. Whether the induction of the system is activated by specific or general signals associated with phage infection or stress needs to be determined. The prophage-associated localization could also contribute to the dissemination of the AbiF module within the bacterial population. This study emphasizes a unique position of AbiF system in the evolutionary path on the crossroad of previously characterized type III TA and ancestral CRISPR-Cas13 sharing either functional or structural similarities. Future comparative studies will decipher the specificity of RNase activity associated with the AbiF toxicity inside the bacterial cell. Overall, our findings further highlight the increasing evidence for the role of RNAs as key components of numerous anti-phage defense systems. By characterizing a prophage-located AbiF-like system, this work is providing insights into the RNA-based bacteria-phage interaction mechanisms paving the way for future biotechnological and health applications.

## Materials and methods

### Bacterial strains and growth conditions

*C. difficile* and *E. coli* strains used in this study are described in Supplementary Data Table S1. *C. difficile* strains were grown in anaerobic conditions (5% H_2_, 5% CO_2_, and 90% N_2_), using an anaerobic chamber (Jacomex) in TY (62) or Brain Heart Infusion (BHI, Difco) medium. Thiamphenicol (Tm, 7.5 µg/mL), cefoxitin (Cfx, 25 µg/mL) and cycloserine (Cs, 250 µg/mL) were added when needed. *E. coli* strains were grown in aerobic conditions in LB (63) with ampicillin (Amp, 100 µg/mL) and chloramphenicol (Cm, 15 µg/mL) when necessary.

### Plasmid construction and conjugation into *C. difficile* strains

All plasmids and primers used in this study are listed in Supplementary Data Table S2 and S3, respectively. All derived plasmids were transformed into *E. coli* NEB10β strain and inserts were verified by sequencing. Then, they were transformed into *E. coli* HB101 (RP4) strain for subsequent transfer by conjugation into *C. difficile* strains. The HB101 donor and *C. difficile* receptor strains were grown overnight in their respective media supplemented with antibiotics when needed. One milliliter of donor strain was centrifuged at 3,500xg for 5 minutes and the supernatant was discarded. Then, 200 µL of *C. difficile* strain was used to gently resuspend the donor strain in anaerobic conditions. For the R20291 strain, the cell suspension was first heated at 50°C for 10 minutes (64). The co-culture was then spotted onto BHI plate and incubated for 8 hours. The cells were then collected with 600µL PBS, plated on BHI supplemented with Cfx, Cs and Tm and incubated for 48 to 72h at 37°C. Transconjugants were verified by PCR for the presence of the plasmid.

### Mutant construction and mutagenesis strategy

For the generation of deletion mutants, we used an allelic exchange method (23) with the toxin from a type I TA system for counter-selection. The pMSR0 plasmid used for this method encodes the toxin *CD2517.1* under the control of the inducible promoter P_tet_ and the antitoxin RCd8 under the control of its own promoter. The homology arms were designed to have 800 to 1,000 bp of upstream and downstream homology to the chromosomal sequence to delete. They were amplified by PCR from the genomic sequence of the *C. difficile* R20291 strain (Supplementary data Table S2 and S3 for plasmid and primers description). Colonies were tested by PCR followed by sequencing to confirm the deletion. RCd22 and phi027 deletion mutants were generated in the *C. difficile* R20291 background and phiCD630-1 was generated in *C. difficile* 630Δ*erm* ΔphiCD630-2 background (23).

### Bioinformatic analyses for distribution and position of AbiF/Abi_2 systems

The presence of proteins assigned to AbiF/Abi_2 COG4823 and pfam07751 family was analyzed in 47,545 complete prokaryotic genome assemblies (chromosome level assembly, November 2023) both from Refseq and Genbank available in NCBI database with their taxonomic description (Table S6, S7). The phylogenetic tree was constructed using iTOL v7 web site including the count of the AbiF/Abi_2 family proteins. The information on the flanking genes located at positions from -−5 to +5 from the *abiF/abi2* gene has been extracted from the genomic data. All proteins were annotated by PSIBLAST 2.16.1 version using clusters of orthologous genes (COG), conserved domain (CD) and PFAM profiles from the conserved domain (CDD) database with E-value equal to 1e-4 and other default parameters settings. Genomes in which the *abiF/abi2* gene was detected were further analyzed with geNomad v1.8.0 (28) to determine if the *abiF/abi2* gene was located on a mobile genetic element (Table S8). The coordinates of the prophages identified with geNomad were compared with the coordinates of the *abiF/abi2* gene to determine which were overlapping (Table S8).

### Phenotypic analysis for growth kinetics and spotted growth assay

*C. difficile* strains were grown overnight in TY broth supplemented with Tm and then inoculated at an OD_600_ of 0.05 in fresh medium with Tm. Growth kinetics were measured manually, using a spectrophometer (Fisher Scientific Cell Density Meter 40), taking OD_600_ every 1 hour during 10 hours. Alternatively, a microplate reader (Cerillo) was used for growth curve analysis. For induction of the P_tet_ promoter of pDIA6103 plasmid derivatives, anhydrotetracycline (ATc, 250 ng/mL) was added at an OD_600_ of 0.4. For drop assay, dilutions (from 10^-1^ to 10^-6^) of an exponential phase grown culture were spotted onto TY agar plates containing Tm (15 µg/mL) and ATc (250 ng/mL).

### Alkaline phosphatase activity assays

*C. difficile* strains containing the *phoZ* fusion plasmids were grown in TY broth supplemented with Tm and harvested at the end of the exponential phase. Samples were stored at −20°C and the alkaline phosphatase assay was performed as previously described (23). Time elapsed for the assay was recorded (Δt in minutes). The absorbance at both OD_420_ and OD_550_ was taken after the alkaline phosphatase reaction. Activity units were calculated and normalized using the following formula: ((OD_420_ – (1.75 × OD_550_)) × 1000)/(Δt × OD_600_ × vol. cells (mL)).

### RNA extraction, qRT-PCR, Northern blot, and 5ʹ/3ʹRACE

For RNA extraction, *C. difficile* strains were grown in TY broth and harvested at various growth phases. *C. difficile* strains carrying pDIA6103 plasmid derivatives were grown in TY with Tm and ATc (250 ng/mL) was added at an OD_600_ of 0.4 for induction of the P_tet_ promoter. Cells were harvested after 4 hours of induction. RNA were extracted using Trizol (Sigma), as previously described (65). Northern Blot and 5’/3’RACE experiments were performed as previously described (4).

### MAPS experiment coupled with mass spectrometry or RNA-sequencing

MAPS experiment coupled with RNA-Sequencing or mass spectrometry was performed and adapted to *C. difficile* from (32). The R20291 ΔRCd22 strain carrying the p117 (pDIA6103 carrying the MS2-tag alone, under the control of the P_tet_ inducible promoter) or p220 plasmid (pDIA6103 carrying MS2-tag and a 6 nucleotides spacer to the 5’-end of RCd22, under the control of the P_tet_ inducible promoter) was grown in 250mL of TY with 7.5µg/mL Tm until the OD_600_ reached 1. Then the expression of the tagged ncRNA RCd22 was induced with 100 ng/mL ATc for 10 min and the culture was cooled on ice. The culture was centrifuged (6,000xg, 15 min, 4°C) and the pellet was resuspended in 5mL ice-cold buffer A (20mM Tris-HCl pH 8, 150mM KCl, 1mM MgCl_2_, 1mM DTT). The mix was transferred into Lysis Matrix B tubes and lysed with a FastPrep machine (30s at setting 6.5, twice with a 2 min pause on ice). Samples were centrifuged at 15,700xg for 15 min at 4°C and the supernatant was transferred into a new tube and maintained at 4°C. For the affinity purification, a poly-prep empty chromatography column was used and all further steps were done at 4°C. First, the column was washed with ultra-pure water (RNase-free) and then 300µL of amylose resin was added. The column was washed with 10mL of buffer A. Then, 1,200pmol of MS2-MBP protein diluted in 6mL of Buffer A was loaded in the column, followed by a wash with 10mL of buffer A. The cell lysate was then loaded onto the column, followed by 3 washes with 10mL of buffer A. The complex was then eluted with 1mL of buffer E (20mM Tris-HCl pH8, 150mM KCl, 1mM MgCl_2_, 1mM DTT, 0.1% Triton X-100, 12mM Maltose). A volume of 1mL of each fraction was used for RNA extraction. One milliliter of phenol was added, mixed vigorously and centrifuged at 16,000xg for 10 min at RT. The upper phase was transferred into a new tube and the organic phase kept for protein precipitation. One milliliter of chloroform/isoamyl alcohol (24/1) was added, followed by a second centrifuge step. The upper phase was transferred into a new tube and 2.5 volumes of 100% cold ethanol and 0.1 volume of 3M sodium acetate pH5.2 were added and the mix was incubated overnight at −20°C. Then, the sample was centrifuged at 4,000xg for 90 min at 4°C, the supernatant was discarded and 500µL of 80% of cold ethanol was added followed by a second centrifugation step at 4,000xg for 45 min at 4°C. The pellet was dried and resuspended in 90µL of ultra-pure water, followed by DNase treatment and inactivation of DNase (TURBO DNA-free, Invitrogen). The RNA was quantified by Qubit and reverse transcription was done on 50 ng of purified RNA with the AMV Reverse Transcriptase (Promega). The resulting cDNA was used as template for PCR to check for the enrichment of MS2-RCd22 in the MAPS eluted fraction. cDNA libraries were prepared using the ScriptSeq kit from Illumina and sequenced by Novogene.

For protein precipitation, 4 volumes of cold acetone were added to the tube containing the organic phase, vortexed and incubated overnight at −20°C. The sample was centrifuged 10 min at 13,000xg and the supernatant was gently discarded. The pellet was dried for 10-30 min and the sample was then processed at the I2BC facility for liquid chromatography coupled to tandem mass spectrometry (LC-MS/MS) analyses. Peak picking, database search against the R20291 proteome, protein inference, and quantitation were done with Mascott (v2.6.2) and experiment-wide grouping with protein cluster analysis were done with Scaffold (v5.0.1) by the Proteomic-Gif (SICaPS) I2BC facilities. We then used the provided “total spectrum counts” as an estimation of the protein quantities and applied the SARtools-DESeq2 (v1.8.1) statistical routines for assessing differential protein enrichment.

### RNA-Sequencing analysis

All codes used during this study are available on github (https://github.com/i2bc/RNAreg_AbiF_CDiff). The RNA sequencing data were processed as described on the github page. Differential expression analysis for enriched genes in MAPS interactomics was performed using the DESeq2 (66) based script SARTools (67), and genes were considered differentially expressed with at least 2 log2 fold change and an adjusted P < 0.05. The analyses were combined in a snakemake pipeline. Briefly, it includes quality control of reads with FastQC and fastQ Screen (and correction with FastP if necessary), creation of an index from the *C. difficile* ribotype R20291 genome, mapping of reads onto the genome with Bowtie2, selection of mapped reads with Samtools and their counting by the FeatureCounts tool of the Subread package on the coding sequences listed in the genome annotation and augmented by the list of ncRNAs selected in a previous analysis (68), as well as a differential gene expression analysis with the SARTools package using the DESeq2 method. All parameters and software versions are defined in an associated configuration file, enabling this pipeline to be re-used for other MAPS analyses involving other ncRNAs and/or genomes. Raw sequencing data have been submitted to ENA with the accession number PRJEB87349.

### Construction, production and protein (co-)purification

The *abiF_Cd_* gene from the *C. difficile* strain R20291 was synthesized and cloned into a pET-16b vector, containing a N-terminal 10X-histidine tag, under the control of the T7*lac* promoter (Supplementary Data Table S1 and S3 for strains and plasmids). The ncRNA RCd22 sequence with the terminator region was amplified from the R20291 strain and cloned into the same plasmid, at a different location, under the control of the T7 promoter (Supplementary data Table S3 for primers). The plasmid was transformed into *E. coli* BL21(DE3) cells and grown overnight at 37°C, 180 RPM in LB medium supplemented with Cm (15 µg/mL) and Amp (100 µg/mL). A sub-culture was then grown at 37°C, 180 RPM until an OD_600_ of 0.7. IPTG was then added to a final concentration of 1mM and cells were incubated at 37°C, 180 RPM during 4h. Cells were harvested by centrifugation at 4,000xrpm g for 10 min at 4°C. Cells were resuspended in lysis buffer (20mM Na_2_HPO4 pH7.5, 1M NaCl, 5% glycerol, 2mM ß-mercaptoethanol, cOmplete mini EDTA-free protease inhibitor cocktail, Roche) and lysed by sonication. The lysate was centrifuged at 4,000xg for 15 min at 4°C, the supernatant was passed through a 0.45µm filter and was loaded on a Ni2+NTA column. The complex AbiF_Cd_-His-tag-RCd22 was eluted by FPLC with AKTA system using elution buffer (Na_2_HPO4 pH7.5, 400mM NaCl, 5% glycerol, 2mM ß-mercaptoethanol, 500mM imidazole). Fractions with the complex eluted were pooled and twofold diluted with dilution buffer (Na_2_HPO4 pH7.5, 5% glycerol, 2mM ß-mercaptoethanol) to decrease the NaCl and imidazole concentration. In an attempt to purify the AbiF_Cd_-His tag protein alone and remove RCd22, benzonase (12.5 U/mL) and MgCl_2_ (10 mM) were added and the sample incubated for 1h with shaking at 180 RPM at room temperature. Then, the sample was loaded on heparin column and AbiF_Cd_-His tag protein was eluted with an AKTA FPLC system using elution buffer (Na_2_HPO4 pH7.5, 2M NaCl, 5% glycerol, 2mM ß-mercaptoethanol). Finally, gel filtration was performed to reduce the NaCl concentration to 200mM and the protein was conserved in 10% glycerol at −80°C. Production of AbiF_Cd_-His tag protein was verified at every step by SDS-PAGE followed by Coomassie staining and Western Blot with an HRP anti-His tag antibody (Proteintech, HRP-66005). To verify ncRNA RCd22 copurification from the affinity column, an RNA extraction was performed directly from the eluted protein purification fraction using phenol-chloroform, following by a gel electrophoresis and RT-qPCR.

### AlphaFold structure prediction

The structural model of AbiF_Cd_ protein homodimer was obtained with AlphaFold2-Multimer (69), using the ColabFold implementation (70) (v1.5.2, commit 3574273 from 24 Feb 2023, which uses AlphaFold version 2.3). First, a multiple sequence alignment (MSA) of AbiF_Cd_ homologs was obtained using MMseqs2 (71) (commit 4148e09, 30 Jan 2023) to query the UniRef30 database version 2202 (72). Then, two concatenated copies of this MSA were used as input to AlphaFold2-Multimer (multimer_v3 parameters) to generate 5 models. The best out of 5 was selected using the AlphaFold2-Multimer combined score (0.8 ipTM + 0.2 pTM). For evolutionary conservation mapping, we used the MSA generated by MMseqs2 and retained only the first 100 sequences (already displaying divergence down to 38% sequence identity with the query AbiF_Cd_ sequence). The structural models of RNA-protein interactions were predicted using the AlphaFold3 web server (73). The best model out of 25 models (5 independent web server runs), ranked by protein-RNA ipTM score while controlling for good overall ipTM and pTM scores, was used for display and analysis.

### Preparation of RNAs for *in vitro* experiments

Transcription of RNAs used in this study (RCd22, *CDR20291_1462, 1558, 2768, 0538, 1829, 3357*) was achieved using PCR products containing the sequence of the RNA downstream of the T7 promoter, introduced along with the primer (Table S3). After purification, these PCR products were used as templates for *in vitro* transcription using T7 RNA polymerase. RNAs were treated with DNase I, purified on an 6% polyacrylamide-8 M urea gel, eluted with 0.5 M ammonium acetate, 0.1 mM EDTA, and 0.1% SDS and precipitated in cold absolute ethanol. RCd22 was labeled with T4 polynucleotide kinase (Fermentas) and [γ^32^P] ATP and ultimately purified on a 6% polyacrylamide-8M urea gel and eluted as described above.

### Electrophoretic mobility shift assay (EMSA)

5’-end radiolabeled RCd22 (10,000 cps/sample, <1pM), and cold mRNAs were denatured separately by incubating at 90°C for 1 min in 100 mM Tris-HCl pH 7.5, 300 mM KCl, 200 mM NH_4_Cl, then cooled down for 1 min on ice and renatured at RT for 10 min after addition of 10 mM MgCl_2_. Complexes were formed at 37°C for 15 min. After the addition of 1 volume of glycerol blue loading buffer, the samples were loaded on a native 6% polyacrylamide gel containing 10 mM MgCl_2_ and migrated at 300 V and 4°C in 1X Tris Borate buffer with 10 mM MgCl_2_ before autoradiography.

## Acknowledgments

This work was supported by grants from the Agence Nationale de la Recherche (the “ADI 2021” project funded by IDEX Paris-Saclay, ANR-11-IDEX-0003-02, M.S. PhD grant; “CloSTARn”, ANR-13-JSV3-0005-01 to O.S., “CdiffRib” ANR-22-CE15-0020-01 to O.S., to I.C. and to C.T.-N., “CDefenseRNA”, ANR-24-CE35-2137-01 to O.S. and to J.A.; “ESPRINet”, ANR-18-CE45-0005-01 to J.A.), the Institut Universitaire de France (to O.S.), the University Paris-Saclay, the Institute for Integrative Biology of the Cell, the DIM-1HEALTH regional Ile-de-France program (LSP grant no. 173403, V.K. PhD fellowship), Natural Sciences and Engineering Research Council of Canada (NSERC discovery grants number RGPIN-2020-05776, to L.-C.F.) and Canadian Institutes of Health Research (CIHR, No.506900, to L.-C. F.). L.-C.F. is a member of the Clinical Research Center of the Centre Hospitalier Universitaire de Sherbrooke (CRCHUS). We are grateful to Dr Muriel Gondry and Carine Tellier for invaluable advices with protein purification procedure and to Pierre Boudry, Anaïs Boutserin and Kimberley Casado for help in initial steps of the project. We thank Benoît Madec for his help in data analysis. This work has benefited from the facilities and expertise of the Proteomic-Gif (SICaPS) platform of I2BC (Institute for Integrative Biology of the Cell, CEA, CNRS, Université Paris-Saclay, Gif-sur-Yvette Cedex, France). We thank members of the SICaPS platform for assistance with mass spectrometry analysis. We thank the BIOI2 platform for making the ColabFold pipeline easily accessible at the I2BC. Computations were made on the supercomputers mp2 managed by the Center for Computational Science at the Université de Sherbrooke.

## Supporting information

**Fig. S1 – Functions enriched in the vicinity of the *abiF/abi_2* gene located in prophages.** (**A**) Functions at the −5 to +5 positions surrounding *abiF/abi_2* located in prophages (N=288 genomes). (**B**) Relative proportion of functions in prophages vs 2,312 genomes where *abiF/abi_2* is not located in a prophage.

**Fig. S2 - Effect of *abiF_Cd_* overexpression on *C. difficile* and *E. coli*.** (**A**) Growth curve and (**B**) CFU/ml of *C. difficile* R20291 carrying p*/abiF_Cd_* or empty plasmid (empty) in TY supplemented with Tm and with (ATc +) or without induction. Induction of the P_tet_ promoter by 250ng/mL ATc is indicated by the red arrow. (**C**) Growth curve of *E. coli* carrying p*/abiF_Cd_* or empty plasmid (empty) in LB supplemented with Cm and 250ng/mL ATc (indicated by a red arrow). Plotted values represent the mean standard error of the mean (N = 3 biologically independent samples).

**Fig. S3 – Growth controls for phenotypic assay in CD156 strains carrying the different plasmid constructions.** (**A**) *C. difficile* CD156 strains carrying empty plasmid (empty), overexpressing *abiF_Cd_* (p*/abiF_Cd_),* or co-expressing *abiF_Cd_* and RCd22 (*in cis* or *in trans,* with or without mutations 1 or 2) were grown in TY supplemented with Tm without induction (ATc -). (**B**) Biological triplicate of spot assay of CD156 strains carrying empty plasmid (1), overexpressing *abiF_Cd_* (2), co-expressing *abiF_Cd_* and RCd22 *in cis* (3), with mutation 1 (4) or mutation 2 (7) or in *trans* (5) with mutation 1 (6) or mutation 2 (8). Spot assay is realized on TY plate supplemented with Tm and with or without inducer (ATc + or -).

**Fig. S4 - Effect of RCd22 deletion on bacterial growth.** (**A**) Relative expression of RCd22 to the reference 16S RNA gene in *C. difficile* R20291 and ΔRCd22 strains. (**B**) Growth curve of *C. difficile* R20291 strain in TY. Plotted values represent the mean standard error of the mean (N = 3 biologically independent samples).

**Fig. S5 – Expression level of *abiF_Cd_* and RCd22 mutated or not in phenotypic assays.** For each strain carrying the different plasmids, the expression of *abiF_Cd_* (**A**), RCd22 (**B**), RCd22mut1 (**C**) and RCd22mut2 (**D**) were controlled by RT-qPCR. Total mRNAs were collected after 2h of induction with ATc, during phenotypic assay. The level of expression is relative to the reference DNApol III gene.

**Fig. S6 – Potential mRNA targets of RCd22 identified by MAPS experiment.** (**A**) Volcano plot of mRNA targeted by RCd22 after MAPS experiment. Red dots indicate enriched RNA targeted by RCd22 compared to the control (MS2tag alone). (**B**) Hybridization energy of RCd22 with five candidate target sequences calculated with IntaRNA. Sequence of mRNA targets and RCd22 started from predicted transcript starting site to predicted terminator. In case of operon, the sequence started from the intergenic region of the target gene. (**C**) Shift assay performed with radioactive RCd22 (RCd22*) transcribed *in vitro* and mixed with different concentrations of potential mRNA targets identified by MAPS and transcribed *in vitro*. mRNA of *abiF_Cd_* (R20291_1462) was also tested.

**Fig. S7 – Copurification of Histag-AbiF_Cd_ protein with RCd22.** (**A**) Verification of *abiF_Cd_*-Histag effect on growth of *E. coli* BL21(DE3) in LB supplemented with Cm, and with or without induction (1mM IPTG, indicated by a red arrow). Plotted values represent the mean standard error of the mean (N = 3 biologically independent samples). (**B**) Coomassie and Western Blot (HRP anti-Histag antibody) of total protein extract from *E. coli* BL21(DE3) carrying the p343 plasmid (Histag-*abiF_Cd_*) before and after 4h of induction with 1mM IPTG. (**C**) Coomassie and Western Blot (HRP anti-Histag antibody) of total protein extract from *E. coli* BL21(DE3) carrying the p353 plasmid (His-tag-*abiF_Cd_* and RCd22*_trans_*) before and after 4h of induction with 1mM IPTG. (**D**) Graph of AbiF_Cd_-His-tag copurification during HisTrap affinity column, obtained by AKTA FPLC system.

**Fig. S8 - Structural alignment of AlphaFold3 model of RCd22-AbiF_Cd_ complex with experimental structure of PbAbiF dimer bound to ncRNA (PDB identifier: 8vz6** (37)). The AlphaFold3 model, colored as in Fig. 8B/C (AbiF_Cd_ in orange and green, RCd22 in purple with *abiF* motifs in cyan), was structurally aligned with the experimental structure of the PbAbiF dimer (in light grey) bound to two copies of a ncRNA (in dark grey). (**A**) Overall view. (**B**) Focus on the region of amino acids R202 and H207 in AbiF_Cd_, structurally aligned to R210 and H215 in PbAbiF.

**Table S1 - Strains used in this study.**

**Table S2 - Plasmids used in this study.**

**Table S3 - Primers used in this study.**

**Table S4 - RNA-seq results from MAPS experiment.**

**Table S5 - Mass spectrometry results from MAPS experiment.**

**Table S6 - Distribution of *abiF/ab*i_2 gene.**

**Table S7 - Environment of *abiF/abi_2* gene.**

**Table S8 - Distribution and environment of *abiF/abi_2* gene in prophage elements.**

## References

1. Lopatina A, Tal N, Sorek R. Abortive Infection: Bacterial Suicide as an Antiviral Immune Strategy. Annual Review of Virology. 2020;7(1):371–84.

2. Rousset F, Sorek R. The evolutionary success of regulated cell death in bacterial immunity. Curr Opin Microbiol. 2023 Aug;74:102312.

3. Georjon H, Bernheim A. The highly diverse antiphage defence systems of bacteria. Nat Rev Microbiol. 2023 Oct;21(10):686–700.

4. Soutourina OA, Monot M, Boudry P, Saujet L, Pichon C, Sismeiro O, et al. Genome-Wide Identification of Regulatory RNAs in the Human Pathogen *Clostridium difficile*. PLOS Genetics. 2013 May 9;9(5):e1003493.

5. Hargreaves KR, Clokie MRJ. *Clostridium difficile* phages: still difficult? Front Microbiol. 2014 Apr 28;5:184.

6. Boudry P, Semenova E, Monot M, Datsenko KA, Lopatina A, Sekulovic O, et al. Function of the CRISPR-Cas System of the Human Pathogen *Clostridium difficile*. mBio. 2015 Sep 1;6(5):e01112–01115.

7. Maikova A, Boudry P, Shiriaeva A, Vasileva A, Boutserin A, Medvedeva S, et al. Protospacer-Adjacent Motif Specificity during *Clostridioides difficile* Type I-B CRISPR-Cas Interference and Adaptation. mBio. 2021 Aug 31;12(4):e0213621.

8. Oliveira PH, Ribis JW, Garrett EM, Trzilova D, Kim A, Sekulovic O, et al. Epigenomic characterization of *Clostridioides difficile* finds a conserved DNA methyltransferase that mediates sporulation and pathogenesis. Nat Microbiol. 2020 Jan;5(1):166–80.

9. Sekulovic O, Fortier LC. Global transcriptional response of *Clostridium difficile* carrying the CD38 prophage. Appl Environ Microbiol. 2015 Feb;81(4):1364–74.

10. Goh S, Ong PF, Song KP, Riley TV, Chang BJ. The complete genome sequence of *Clostridium difficile* phage ϕC2 and comparisons to ϕCD119 and inducible prophages of CD630. Microbiology. 2007;153(3):676–85.

11. Whittle MJ, Bilverstone TW, van Esveld RJ, Lücke AC, Lister MM, Kuehne SA, et al. A Novel Bacteriophage with Broad Host Range against *Clostridioides difficile* Ribotype 078 Supports SlpA as the Likely Phage Receptor. Microbiology Spectrum. 2022 Feb 2;10(1):e02295–21.

12. Garvey P, Fitzgerald GF, Hill C. Cloning and DNA sequence analysis of two abortive infection phage resistance determinants from the lactococcal plasmid pNP40. Appl Environ Microbiol. 1995 Dec;61(12):4321–8.

13. Grafakou A, Mosterd C, Beck MH, Kelleher P, McDonnell B, de Waal PP, et al. Discovery of antiphage systems in the lactococcal plasmidome. Nucleic Acids Research. 2024 Sep 9;52(16):9760–76.

14. Bidnenko E, Chopin MC, Ehrlich SD, Anba J. *Lactococcus lactis* AbiD1 abortive infection efficiency is drastically increased by a phage protein. FEMS Microbiology Letters. 2002 Sep;214(2):283–7.

15. Bidnenko E, Chopin A, Ehrlich SD, Chopin MC. Activation of mRNA translation by phage protein and low temperature: the case of *Lactococcus lactis* abortive infection system AbiD1. BMC Molecular Biology. 2009 Jan 27;10(1):4.

16. Anba J, Bidnenko E, Hillier A, Ehrlich D, Chopin MC. Characterization of the lactococcal *abiD1* gene coding for phage abortive infection. J Bacteriol. 1995 Jul 1;177(13):3818–23.

17. Fineran PC, Blower TR, Foulds IJ, Humphreys DP, Lilley KS, Salmond GPC. The phage abortive infection system, ToxIN, functions as a protein–RNA toxin–antitoxin pair. Proc Natl Acad Sci U S A. 2009 Jan 20;106(3):894–9.

18. Blower TR, Pei XY, Short FL, Fineran PC, Humphreys DP, Luisi BF, et al. A processed noncoding RNA regulates an altruistic bacterial antiviral system. Nat Struct Mol Biol. 2011 Feb;18(2):185–90.

19. Guegler CK, Laub MT. Shutoff of host transcription triggers a toxin-antitoxin system to cleave phage RNA and abort infection. Mol Cell. 2021 Jun 3;81(11):2361–2373.e9.

20. Banh DV, Roberts CG, Morales-Amador A, Berryhill BA, Chaudhry W, Levin BR, et al. Bacterial cGAS senses a viral RNA to initiate immunity. Nature. 2023 Nov;623(7989):1001–8.

21. Bobonis J, Mitosch K, Mateus A, Karcher N, Kritikos G, Selkrig J, et al. Bacterial retrons encode phage-defending tripartite toxin-antitoxin systems. Nature. 2022 Sep;609(7925):144–50.

22. Millman A, Bernheim A, Stokar-Avihail A, Fedorenko T, Voichek M, Leavitt A, et al. Bacterial Retrons Function In Anti-Phage Defense. Cell. 2020 Dec 10;183(6):1551–1561.e12.

23. Peltier J, Hamiot A, Garneau JR, Boudry P, Maikova A, Hajnsdorf E, et al. Type I toxin-antitoxin systems contribute to the maintenance of mobile genetic elements in *Clostridioides difficile*. Commun Biol. 2020 Nov 27;3(1):718.

24. Stabler RA, He M, Dawson L, Martin M, Valiente E, Corton C, et al. Comparative genome and phenotypic analysis of *Clostridium difficile* 027 strains provides insight into the evolution of a hypervirulent bacterium. Genome Biol. 2009;10(9):R102.

25. He M, Miyajima F, Roberts P, Ellison L, Pickard DJ, Martin MJ, et al. Emergence and global spread of epidemic healthcare-associated *Clostridium difficile*. Nat Genet. 2013 Jan;45(1):109–13.

26. Kuntová L, Mašlaňová I, Obořilová R, Šimečková H, Finstrlová A, Bárdy P, et al. *Staphylococcus aureus* Prophage-Encoded Protein Causes Abortive Infection and Provides Population Immunity against Kayviruses. mBio. 2023 Apr 25;14(2):e0249022.

27. Gentile GM, Wetzel KS, Dedrick RM, Montgomery MT, Garlena RA, Jacobs-Sera D, et al. More Evidence of Collusion: a New Prophage-Mediated Viral Defense System Encoded by Mycobacteriophage Sbash. mBio. 2019 Mar 19;10(2):e00196–19.

28. Camargo AP, Roux S, Schulz F, Babinski M, Xu Y, Hu B, et al. Identification of mobile genetic elements with geNomad. Nat Biotechnol. 2024 Aug;42(8):1303–12.

29. Weinberg Z, Lünse CE, Corbino KA, Ames TD, Nelson JW, Roth A, et al. Detection of 224 candidate structured RNAs by comparative analysis of specific subsets of intergenic regions. Nucleic Acids Res. 2017 Oct 13;45(18):10811–23.

30. Koga M, Otsuka Y, Lemire S, Yonesaki T. *Escherichia coli rnlA* and *rnlB* Compose a Novel Toxin–Antitoxin System. Genetics. 2011 Jan;187(1):123–30.

31. Slavcev RA, Hayes S. Blocking the T4 lysis inhibition phenotype. Gene. 2003 Dec 4;321:163–71.

32. Lalaouna D, Prévost K, Eyraud A, Massé E. Identification of unknown RNA partners using MAPS. Methods. 2017 Mar 15;117:28–34.

33. Bidnenko E, Ehrlich D, Chopin MC. Phage operon involved in sensitivity to the *Lactococcus lactis* abortive infection mechanism AbiD1. Journal of Bacteriology. 1995 Jul;177(13):3824–9.

34. Yoon PH, Zhang Z, Loi KJ, Adler BA, Lahiri A, Vohra K, et al. Structure-guided discovery of ancestral CRISPR-Cas13 ribonucleases. Science. 2024 Aug 2;385(6708):538–43.

35. Pillon MC, Gordon J, Frazier MN, Stanley RE. HEPN RNases - an emerging class of functionally distinct RNA processing and degradation enzymes. Crit Rev Biochem Mol Biol. 2021 Feb;56(1):88–108.

36. Anantharaman V, Makarova KS, Burroughs AM, Koonin EV, Aravind L. Comprehensive analysis of the HEPN superfamily: identification of novel roles in intra-genomic conflicts, defense, pathogenesis and RNA processing. Biology Direct. 2013 Jun 15;8(1):15.

37. Zilberzwige-Tal S, Altae-Tran H, Kannan S, Wilkinson ME, Vo SCDT, Strebinger D, et al. Reprogrammable RNA-targeting CRISPR systems evolved from RNA toxin-antitoxins. Cell. 2025 Feb;S0092867425001035.

38. Ainsworth S, Stockdale S, Bottacini F, Mahony J, van Sinderen D. The *Lactococcus lactis* plasmidome: much learnt, yet still lots to discover. FEMS Microbiol Rev. 2014 Sep;38(5):1066–88.

39. Ventura M, Zomer A, Canchaya C, O’Connell-Motherway M, Kuipers O, Turroni F, et al. Comparative Analyses of Prophage-Like Elements Present in Two *Lactococcus lactis* Strains. Applied and Environmental Microbiology. 2007 Dec;73(23):7771–80.

40. Dedrick RM, Jacobs-Sera D, Bustamante CAG, Garlena RA, Mavrich TN, Pope WH, et al. Prophage-mediated defence against viral attack and viral counter-defence. Nat Microbiol. 2017 Jan 9;2:16251.

41. Hayes F. Toxins-antitoxins: plasmid maintenance, programmed cell death, and cell cycle arrest. Science. 2003 Sep 12;301(5639):1496–9.

42. Durand S, Jahn N, Condon C, Brantl S. Type I toxin-antitoxin systems in *Bacillus subtilis*. RNA Biol. 2012 Dec;9(12):1491–7.

43. Meeske AJ, Nakandakari-Higa S, Marraffini LA. Cas13-induced cellular dormancy prevents the rise of CRISPR-resistant bacteriophage. Nature. 2019 Jun;570(7760):241–5.

44. Makarova KS, Wolf YI, Iranzo J, Shmakov SA, Alkhnbashi OS, Brouns SJJ, et al. Evolutionary classification of CRISPR-Cas systems: a burst of class 2 and derived variants. Nat Rev Microbiol. 2020 Feb;18(2):67–83.

45. Jain I, Kolesnik M, Kuznedelov K, Minakhin L, Morozova N, Shiriaeva A, et al. tRNA anticodon cleavage by target-activated CRISPR-Cas13a effector. Sci Adv. 2024 Apr 26;10(17):eadl0164.

46. Jurėnas D, Fraikin N, Goormaghtigh F, Van Melderen L. Biology and evolution of bacterial toxin-antitoxin systems. Nat Rev Microbiol. 2022 Jun;20(6):335–50.

47. Saunier M, Fortier LC, Soutourina O. RNA-based regulation in bacteria-phage interactions. Anaerobe. 2024 Jun;87:102851.

48. Mayo-Muñoz D, Li H, Mestre MR, Pinilla-Redondo R. The role of noncoding RNAs in bacterial immunity. Trends in Microbiology. 2025 Feb 1;33(2):208–22.

49. Carrier MC, Lalaouna D, Massé E. A game of tag: MAPS catches up on RNA interactomes. RNA Biology. 2016 May 3;13(5):473–6.

50. Tomasini A, Moreau K, Chicher J, Geissmann T, Vandenesch F, Romby P, et al. The RNA targetome of *Staphylococcus aureus* non-coding RNA RsaA: impact on cell surface properties and defense mechanisms. Nucleic Acids Res. 2017 Jun 20;45(11):6746–60.

51. Lalaouna D, Desgranges E, Caldelari I, Marzi S. MS2-Affinity Purification Coupled With RNA Sequencing Approach in the Human Pathogen *Staphylococcus aureus*. Methods Enzymol. 2018;612:393–411.

52. Manikandan P, Nadig K, Singh M. Large-scale Purification of Type III Toxin-antitoxin Ribonucleoprotein Complex and its Components from *Escherichia coli* for Biophysical Studies. Bio Protoc. 2023 Jul 5;13(13):e4763.

53. Short FL, Pei XY, Blower TR, Ong SL, Fineran PC, Luisi BF, et al. Selectivity and self-assembly in the control of a bacterial toxin by an antitoxic noncoding RNA pseudoknot. Proceedings of the National Academy of Sciences. 2013 Jan 15;110(3):E241–9.

54. Short FL, Akusobi C, Broadhurst WR, Salmond GPC. The bacterial Type III toxin-antitoxin system, ToxIN, is a dynamic protein-RNA complex with stability-dependent antiviral abortive infection activity. Sci Rep. 2018 Jan 17;8(1):1013.

55. Manikandan P, Sandhya S, Nadig K, Paul S, Srinivasan N, Rothweiler U, et al. Identification, functional characterization, assembly and structure of ToxIN type III toxin-antitoxin complex from *E. coli*. Nucleic Acids Res. 2022 Feb 22;50(3):1687–700.

56. Blower TR, Short FL, Rao F, Mizuguchi K, Pei XY, Fineran PC, et al. Identification and classification of bacterial Type III toxin-antitoxin systems encoded in chromosomal and plasmid genomes. Nucleic Acids Res. 2012 Jul;40(13):6158–73.

57. Samson JE, Bélanger M, Moineau S. Effect of the Abortive Infection Mechanism and Type III Toxin/Antitoxin System AbiQ on the Lytic Cycle of *Lactococcus lactis* Phages. J Bacteriol. 2013 Sep;195(17):3947–56.

58. Rao F, Short FL, Voss JE, Blower TR, Orme AL, Whittaker TE, et al. Co-evolution of quaternary organization and novel RNA tertiary interactions revealed in the crystal structure of a bacterial protein-RNA toxin-antitoxin system. Nucleic Acids Res. 2015 Oct 30;43(19):9529–40.

59. Goeders N, Chai R, Chen B, Day A, Salmond GPC. Structure, Evolution, and Functions of Bacterial Type III Toxin-Antitoxin Systems. Toxins (Basel). 2016 Sep 28;8(10):282.

60. Blower TR, Fineran PC, Johnson MJ, Toth IK, Humphreys DP, Salmond GPC. Mutagenesis and Functional Characterization of the RNA and Protein Components of the toxIN Abortive Infection and Toxin-Antitoxin Locus of *Erwinia*. Journal of Bacteriology. 2009 Oct;191(19):6029–39.

61. Bélanger M, Moineau S. Mutational Analysis of the Antitoxin in the Lactococcal Type III Toxin-Antitoxin System AbiQ. Appl Environ Microbiol. 2015 Jun;81(11):3848–55.

62. Dupuy B, Sonenshein AL. Regulated transcription of *Clostridium difficile* toxin genes. Mol Microbiol. 1998 Jan;27(1):107–20.

63. Bertani G. Studies on lysogenesis. I. The mode of phage liberation by lysogenic *Escherichia coli*. J Bacteriol. 1951 Sep;62(3):293–300.

64. Kirk JA, Fagan RP. Heat shock increases conjugation efficiency in *Clostridium difficile*. Anaerobe. 2016 Dec;42:1–5.

65. Peltier J, Soutourina O. Identification of c-di-GMP-Responsive Riboswitches. Methods Mol Biol. 2017;1657:377–402.

66. Love MI, Huber W, Anders S. Moderated estimation of fold change and dispersion for RNA-seq data with DESeq2. Genome Biology. 2014 Dec 5;15(12):550.

67. Varet H, Brillet-Guéguen L, Coppée JY, Dillies MA. SARTools: A DESeq2- and EdgeR-Based R Pipeline for Comprehensive Differential Analysis of RNA-Seq Data. PLOS ONE. 2016 Jun 9;11(6):e0157022.

68. Kreis V, Toffano-Nioche C, Denève-Larrazet C, Marvaud JC, Garneau JR, Dumont F, et al. Dual RNA-seq study of the dynamics of coding and non-coding RNA expression during *Clostridioides difficile* infection in a mouse model. mSystems. 2024 Nov 27;9(12):e00863–24.

69. Jumper J, Evans R, Pritzel A, Green T, Figurnov M, Ronneberger O, et al. Highly accurate protein structure prediction with AlphaFold. Nature. 2021 Aug;596(7873):583–9.

70. Mirdita M, Schütze K, Moriwaki Y, Heo L, Ovchinnikov S, Steinegger M. ColabFold: making protein folding accessible to all. Nat Methods. 2022 Jun;19(6):679–82.

71. Mirdita M, Steinegger M, Söding J. MMseqs2 desktop and local web server app for fast, interactive sequence searches. Bioinformatics. 2019 Aug 15;35(16):2856–8.

72. Mirdita M, von den Driesch L, Galiez C, Martin MJ, Söding J, Steinegger M. Uniclust databases of clustered and deeply annotated protein sequences and alignments. Nucleic Acids Res. 2017 Jan 4;45(D1):D170–6.

73. Abramson J, Adler J, Dunger J, Evans R, Green T, Pritzel A, et al. Accurate structure prediction of biomolecular interactions with AlphaFold 3. Nature. 2024 Jun;630(8016):493–500.

