## Supplementary material for "Deciphering the RNA-based regulation mechanism of the phage-encoded AbiF system in *Clostridioides difficile*": Figure S1

A

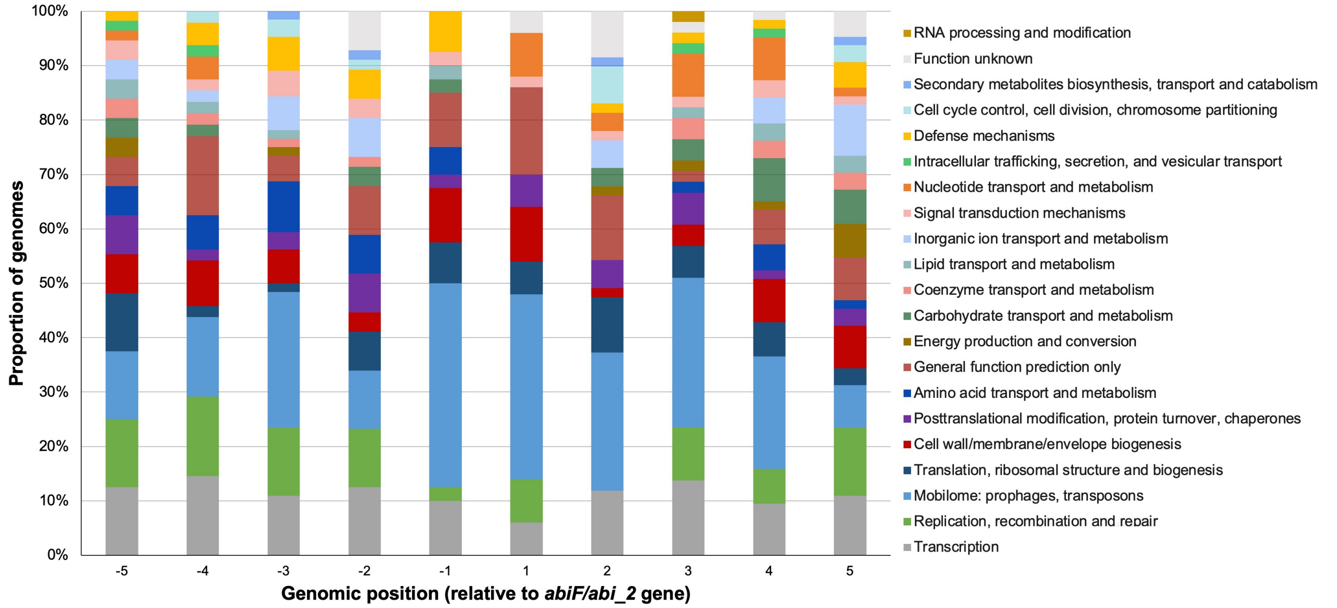


B

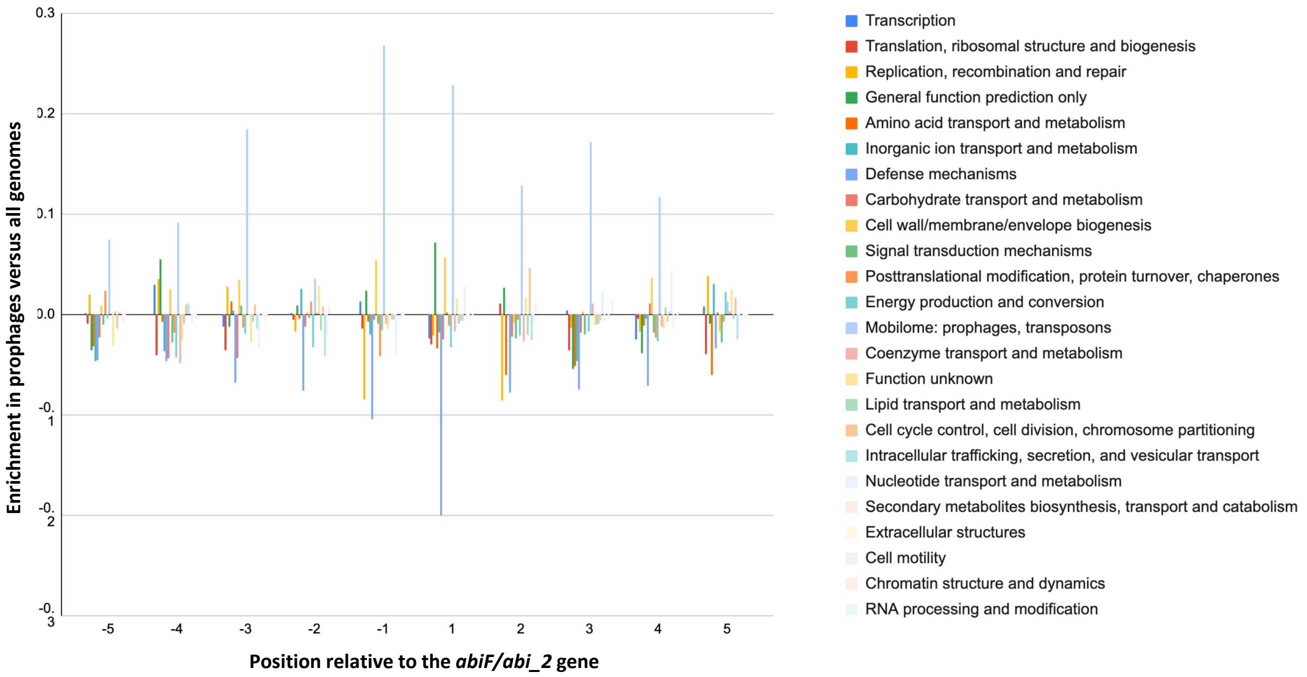


**Fig. S1 – Functions enriched in the vicinity of the *abiF/abi_2* gene located in prophages.**

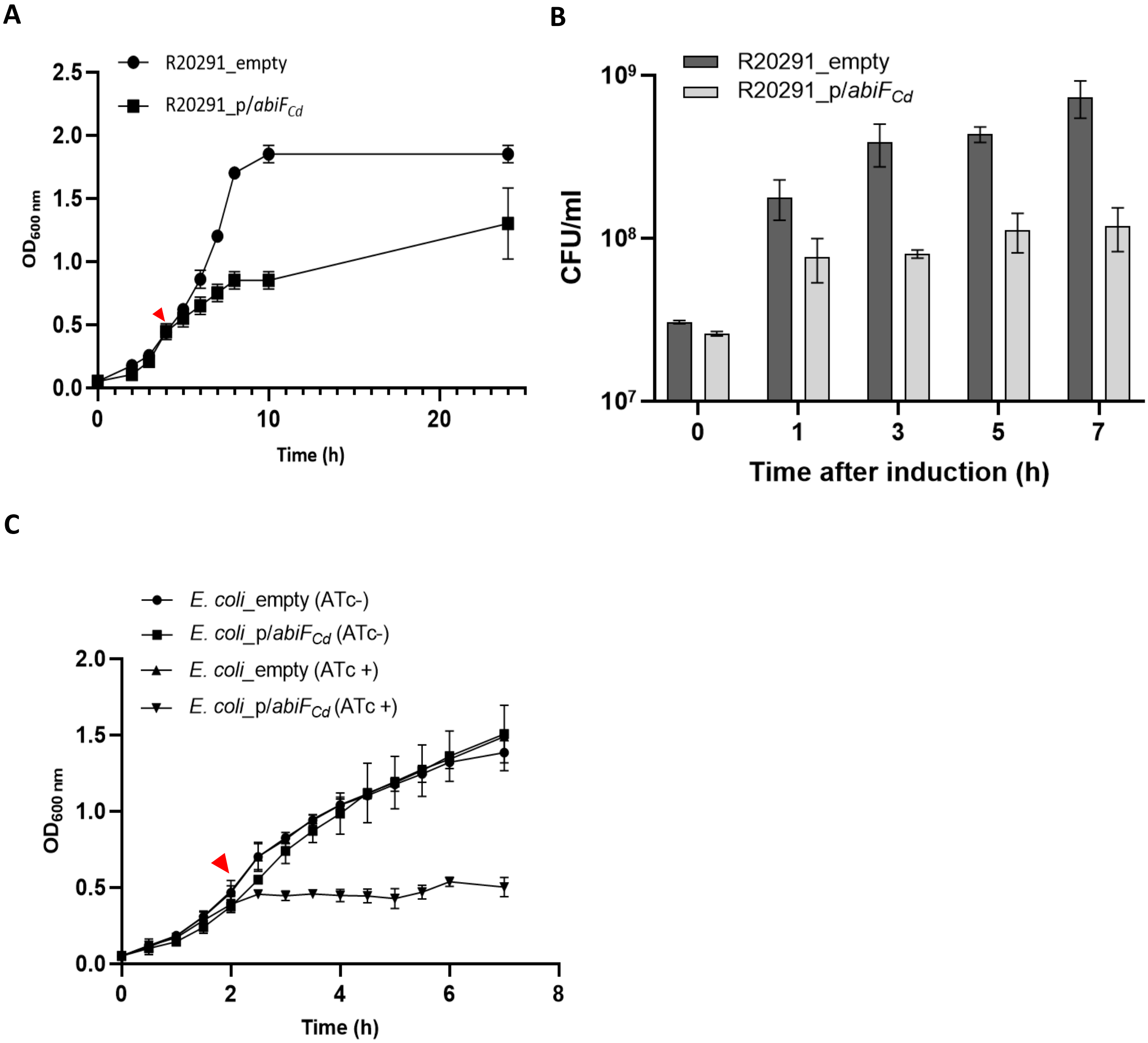


**Fig. S2 - Effect of *abiF*$_{Cd}$ overexpression on *C. difficile* and *E. coli.***

**A.**

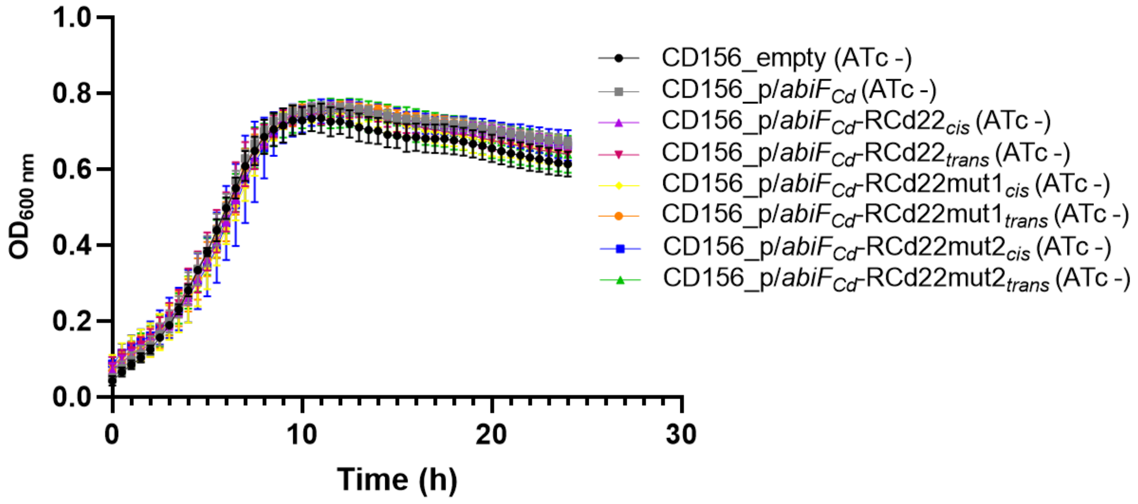


**B.**

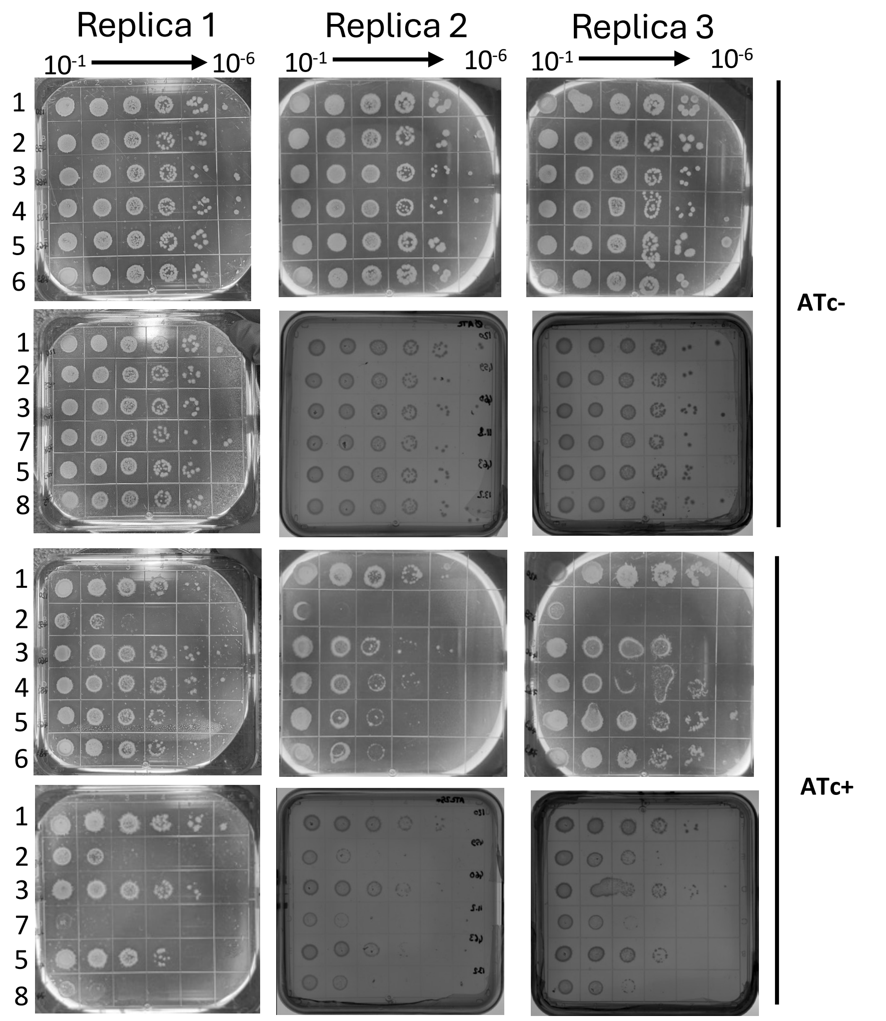


**Fig. S3 - Growth controls for phenotypic assay in CD156 strains carrying the different plasmid constructions.**

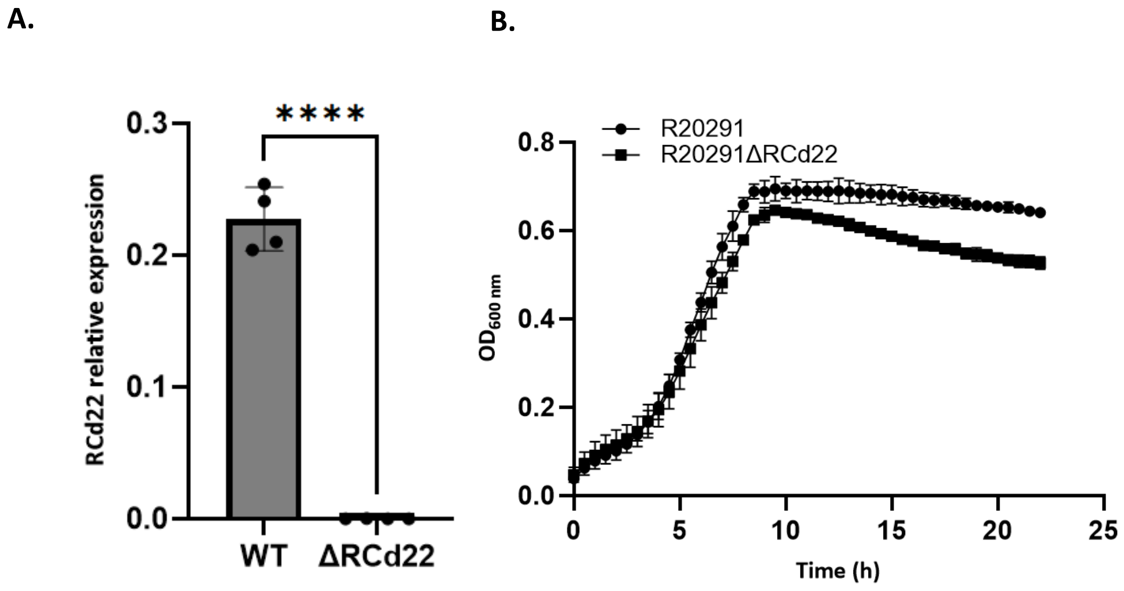


**Fig. S4 – Effect of RCd22 deletion on bacterial growth.**

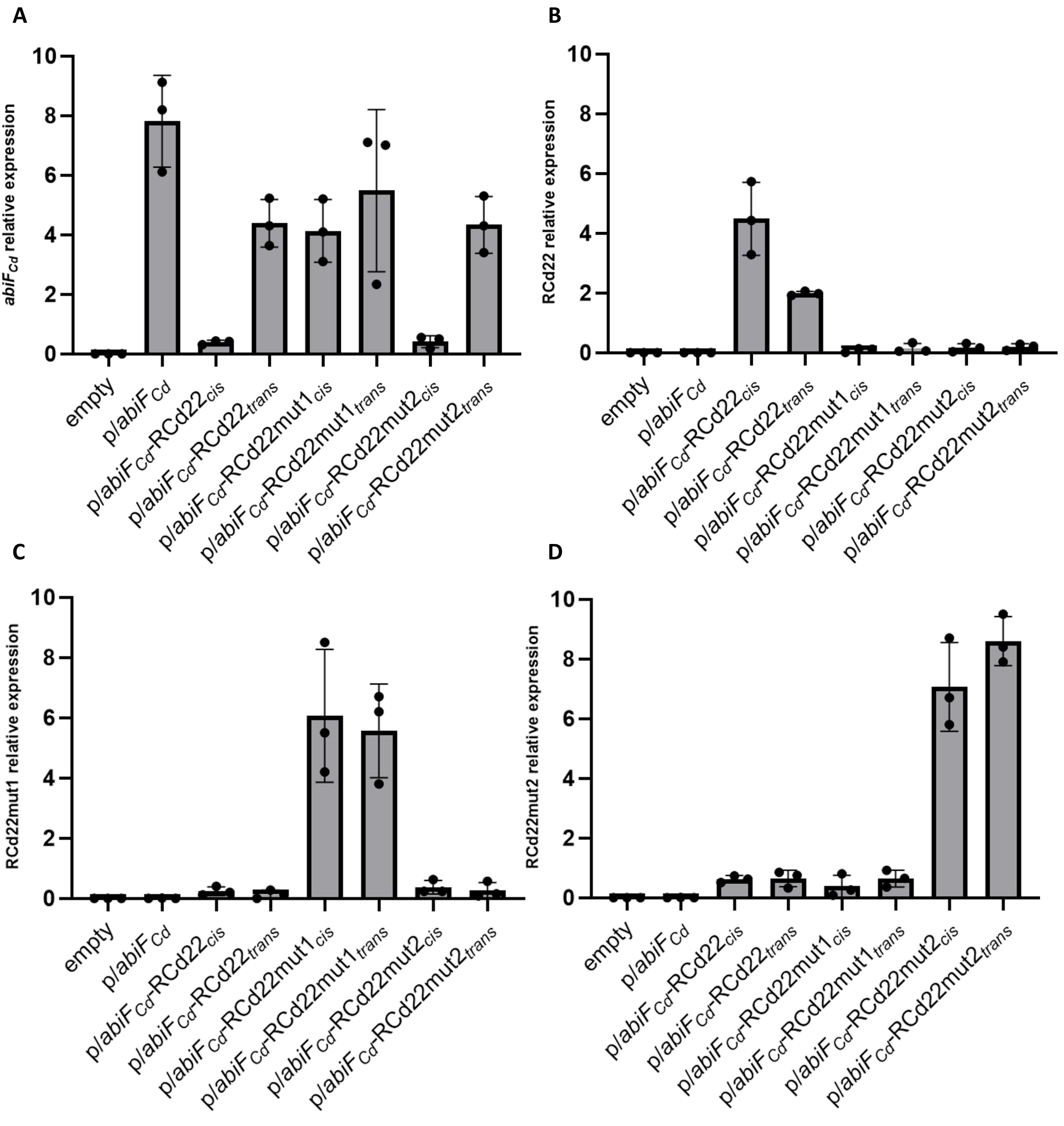


**Fig. S5 - Expression level of *abiF*$_{Cd}$ and RCd22 mutated or not in phenotypic assays.**

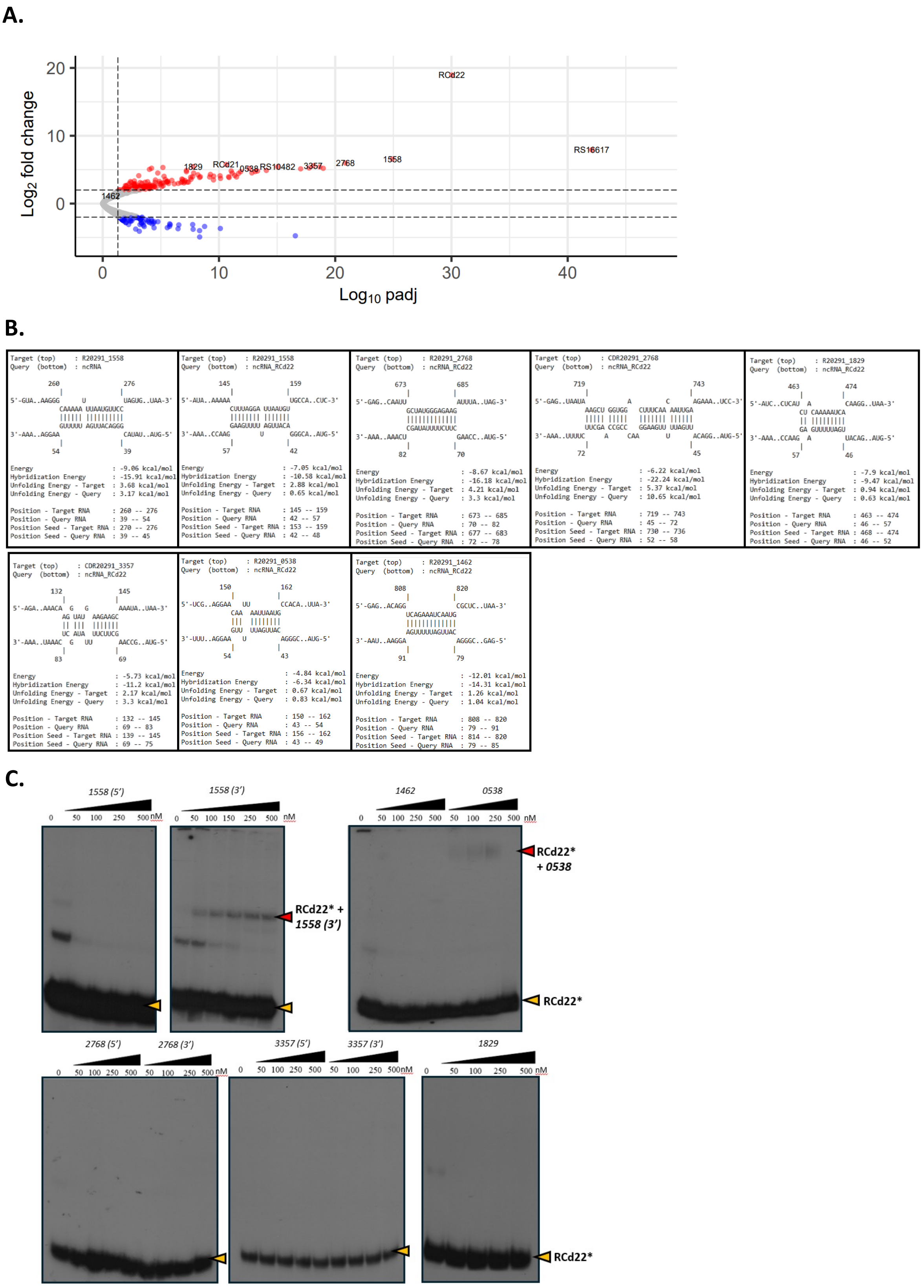


**Fig. S6 - Potential mRNA targets of RCd22 identified by MAPS experiment.**

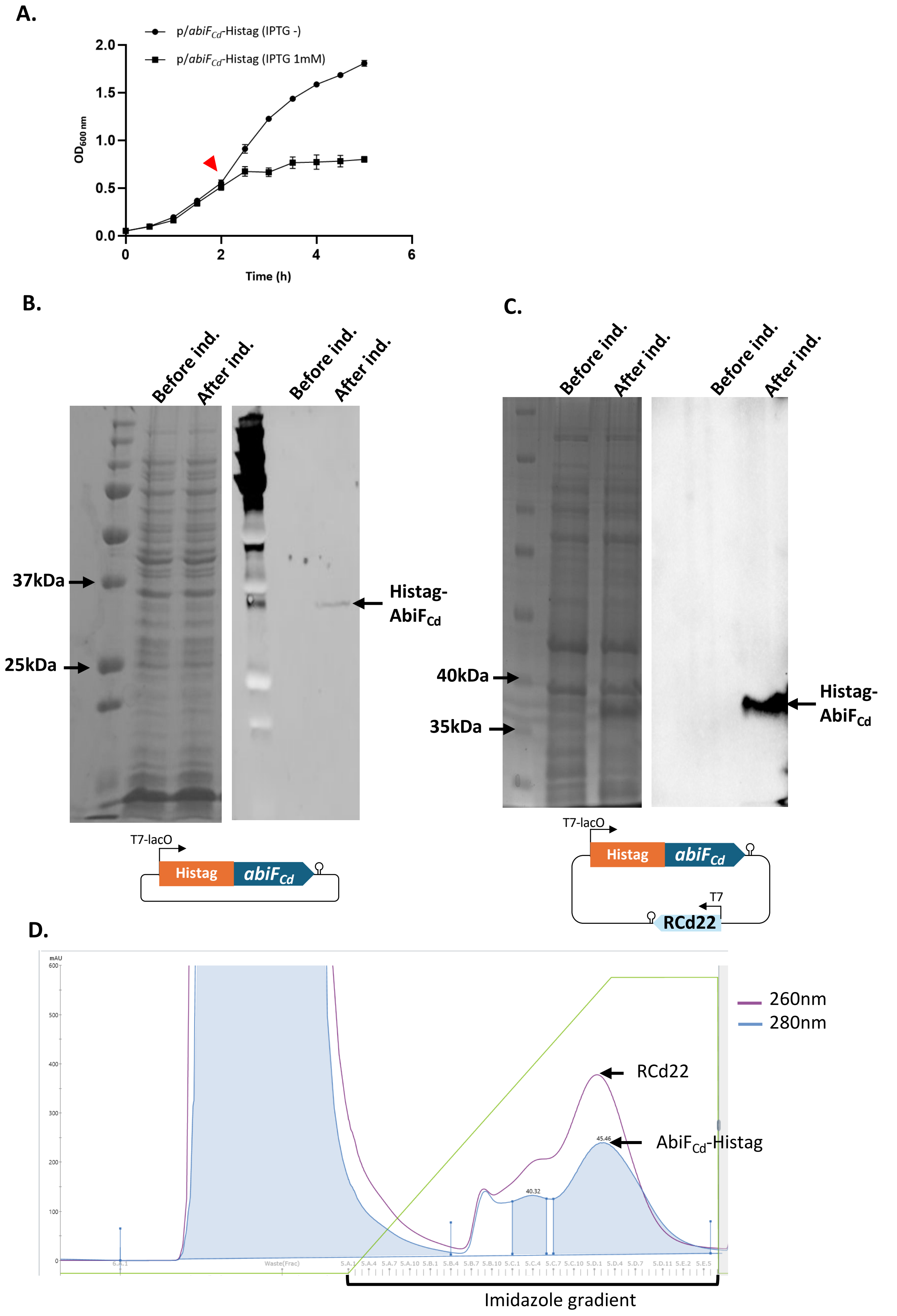


**Fig. S7 – Copurification of Histag-AbiF$_{Cd}$ protein with RCd22.**

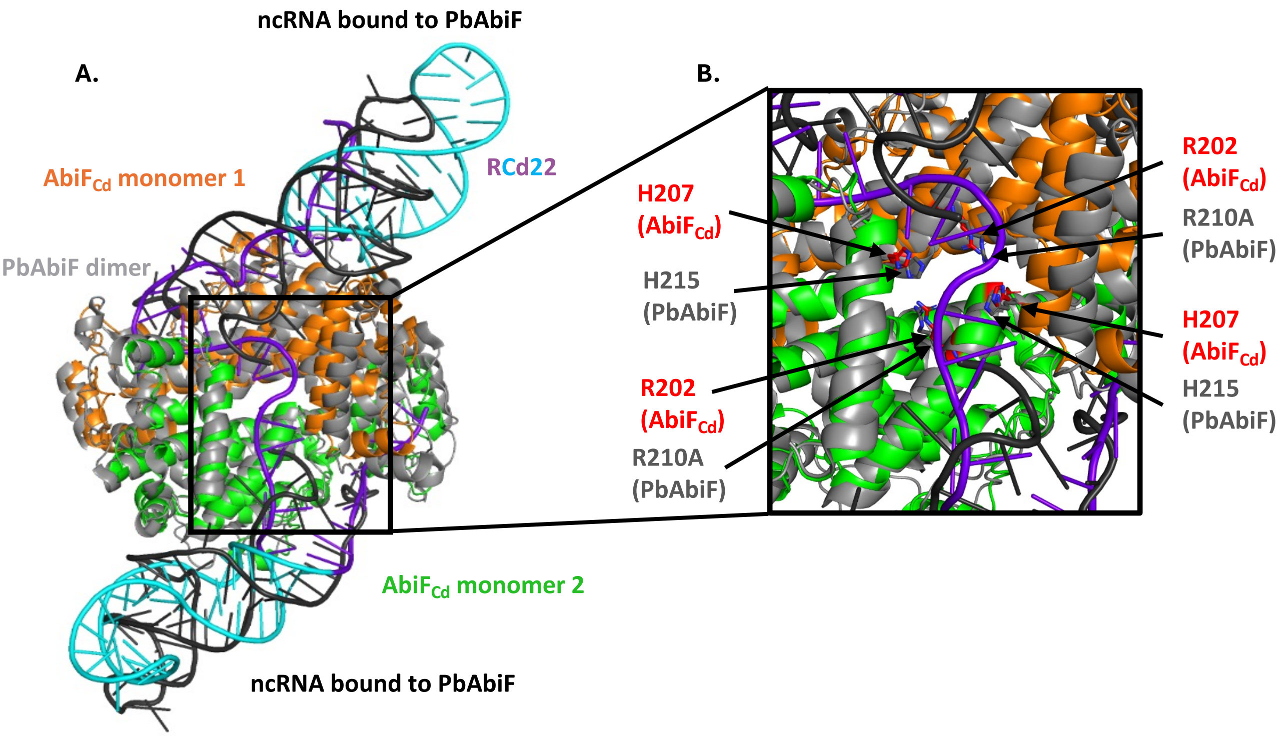


**Fig. S8 - Structural alignment of AlphaFold3 model of RCd22-AbiF$_{Cd}$ complex with experimental structure of PbAbiF dimer bound to ncRNA (PDB identifier: 8vz6).**
